# Membrane Contact Sites Between Chloroplasts and Pathogen Interface Underpin Plant Focal Immune Responses

**DOI:** 10.1101/2021.10.08.463641

**Authors:** Enoch Lok Him Yuen, Zachary Savage, Vanda Adamkova, Cristina Vuolo, Yuanyang Zhou, Yasin Tumtas, Jessica Lee Erickson, Jennifer Prautsch, Andrada I. Balmez, Johannes Stuttmann, Cian Duggan, Francesco Rivetti, Camilla Molinari, Martin Schattat, Tolga O. Bozkurt

## Abstract

Communication between cellular organelles is essential for mounting effective innate immune responses to eliminate pathogens. In plants, the transport of cellular organelles to pathogen penetration sites and their assembly around the host membrane delineating plant-pathogen interface are well-documented. However, whether organelles associate with these specialized plant-pathogen membrane interfaces and the extent to which this process contributes to immunity remain unknown. Here, we discovered defense-related membrane contact sites (MCS) comprising a membrane tethering complex between chloroplasts and the extrahaustorial membrane (EHM) surrounding the pathogen haustorium. The assembly of this membrane tethering complex relies on the association between the chloroplast outer envelope protein CHLOROPLAST UNUSUAL POSITIONING 1 (CHUP1), and its plasma membrane-associated partner, KINESIN-LIKE PROTEIN FOR ACTIN-BASED CHLOROPLAST MOVEMENT 1 (KAC1). Our biochemical assays revealed that CHUP1 and KAC1 interact, while infection cell biology demonstrated their co-accumulation in foci where chloroplasts contact the EHM. Genetic depletion of CHUP1 or KAC1 reduces the deposition of callose—a cell wall material typically deployed to fortify pathogen penetration resistance—around the haustorium, without affecting other core immune processes. Our findings suggest that the chloroplast-EHM attachment complex positively regulates plant focal immunity, revealing the key components and their potential roles in the targeted deposition of defense components at the pathogen interface. These results advance our understanding of organelle-mediated immune responses and highlight the significance of MCS in plant-pathogen interactions.

## Introduction

Filamentous pathogens such as oomycetes and fungi intimately interact with plant hosts, often through specialized infection structures that penetrate the host cells. In response, the invaded plant cell activates a cell-autonomous defense mechanism interface known as focal immunity^1–4^. This defense involves significant cellular reorganization, including organelle relocation, cell-wall reinforcements at pathogen contact sites via callose deposition, and the polarized secretion of antimicrobials^3,5–7^. In addition to the secretory system and the nucleus, organelles such as chloroplasts, mitochondria and peroxisomes accumulate around host cell penetration sites of fungal and oomycete pathogens^5,7,8^. However, the physical association of organelles with the plant-pathogen interface and the extent to which these processes contribute to immunity remain unclear.

Cellular homeostasis relies on efficient interorganelle communication, a process facilitated by long-range vesicle trafficking and short-range membrane contact sites (MCS). MCS are specialized regions where organelles come into close proximity, allowing for the direct transfer of lipids, proteins, signaling molecules, and metabolites. These sites serve as critical gateways for rapid and effective intracellular communication, enabling cells to quickly adapt and respond to stress conditions^9–11^. Recent research has highlighted the emerging roles of MCS in mammalian innate immunity^12^. Moreover, pathogens have developed strategies to target and subvert MCS, perturbing interorganelle communications to undermine host immune defenses and exploit cellular resources^13,14^. However, the involvement of MCS in plant-pathogen interactions remains unexplored. Additionally, the identity and roles of protein-tethering complexes that regulate the functions of MCS in plants are still largely elusive^11^. Understanding these mechanisms could reveal new insights into how organelle communication contributes to immune responses.

Accumulating evidence points to key roles of chloroplasts in the deployment of various plant immune responses^15,16^. Upon immune activation and signaling mediated through mitogen-activated protein kinases (MAPKs), chloroplasts terminate photosynthesis and activate a range of defense responses such as the production of reactive oxygen species (ROS) and defense hormones^17^. During the immune response, chloroplasts alter their morphology by extending stroma-filled tubules, called stromules, that can make contacts with other membranes and organelles such as the nucleus^7,18,19^. In addition, chloroplasts cluster around the nucleus during immune stimulation^20^, a response that is presumed to contribute towards plant defense, possibly through facilitating more efficient chloroplast-to-nucleus signaling. Consistent with the emerging roles of chloroplast in immunity, an increasing number of effectors secreted by pathogens have been found to target chloroplast functions, whereas plants monitor these potential threats using nucleotide-binding leucine-rich repeat (NLR) immune receptors^21–27^.

The Irish potato famine pathogen, *Phytophthora infestans*, can penetrate host cells via specialized infection structures called haustoria, which mediate the delivery of effector proteins inside the host cells. Haustoria are excluded from the host cytoplasm through a newly synthesized plant-derived membrane called the extrahaustorial membrane (EHM)^28–30^. Remarkably, chloroplasts frequently gather around *P. infestans* haustoria and appear to establish tight contacts with the EHM^7^. Given the range of antimicrobial and defense components produced by chloroplasts, their positioning at the pathogen interface likely contributes to focal plant immune responses. The mechanisms underlying chloroplast photorelocation in response to light intensity, involving membrane attachment, detachment, and movement, are relatively well understood, with key components such as CHLOROPLAST UNUSUAL POSITIONING 1 (CHUP1) and KINESIN-LIKE PROTEIN FOR ACTIN-BASED CHLOROPLAST MOVEMENT 1 (KAC1) identified^31–34^. Both CHUP1 and KAC1 are involved in the photorelocation movement of chloroplasts and the association of chloroplasts with the plasma membrane (PM) by regulating short actin filaments on the chloroplast envelope (cp-actin filaments)^31,32,34,35^. However, the extent to which chloroplast movement and positioning around the haustoria contributes to plant immunity remains to be elucidated.

In this study, we investigated the role of the chloroplast movement and membrane anchoring protein CHUP1 in plant immunity against *P. infestans*. Using virus-induced gene silencing (VIGS) and CRISPR knockouts, we found that CHUP1-deficient *N. benthamiana* plants exhibit significantly increased pathogen growth, indicating that CHUP1 positively contributes to immunity. Despite no significant changes in chloroplast movement towards haustoria or other core immune processes such as MAPK-triggered signaling and hypersensitive response (HR) cell death, CHUP1-knockout plants showed reduced callose deposition at haustoria penetration sites, highlighting the involvement of CHUP1 in focal immune responses. We discovered that CHUP1 interacts with the kinesin-like protein KAC1, and both proteins co-accumulate at chloroplast-EHM MCS. This suggests their cooperative role in anchoring chloroplasts to the pathogen interface and enhancing immunity. Consistently, KAC1 accumulated at chloroplast-EHM MCS in a CHUP1-dependent manner, and depletion of KAC1 led to reduced callose deposition around the haustorium and enhanced disease susceptibility. These findings underscore the importance of CHUP1 and KAC1 cooperation in coordinating the tethering of chloroplasts to the pathogen interface and mounting effective immune responses via MCS.

## Results

### CHUP1 positively contributes to immunity against *Phytophthora infestans*

The role of chloroplasts in providing biochemical defense against pathogens is well established^15,16,18^. An emerging yet less understood aspect of chloroplast immunity involves the positioning and movement of epidermal chloroplasts during infection and their contribution to the immune response^20,36^. We previously demonstrated that during infection by the oomycete pathogen *Phytophthora infestans*, epidermal chloroplasts in the model solanaceous plant *Nicotiana benthamiana* accumulate around the haustoria^7^. To investigate this phenomenon further, we performed infection assays on *N. benthamiana* following the downregulation of the chloroplast movement and anchoring gene, *CHUP1*. Silencing the two identified *CHUP1* alleles (*NbCHUP1a* and *NbCHUP1b*) in transgenic *N. benthamiana* expressing GFP in the chloroplast stroma (CP plants) via virus-induced gene silencing (VIGS) (Figure S1A) resulted in significantly higher levels of hyphal growth of tdTomato-expressing *P. infestans* compared to the silencing control (Figures S1B and S1C). To further validate these findings, we generated CRISPR knockout lines lacking *NbCHUP1a* and *NbCHUP1b* (referred to as *chup1* KO plants) from a transgenic parental line of *N. benthamiana* expressing the chimeric protein FNR:eGFP, which targets the plastid stroma (referred to as FNR plants) (Figure S1D). *chup1* KO plants did not exhibit any major developmental defects compared to FNR control plants (Figure S1E). Following infection with *P. infestans*, the mean hyphal growth of the pathogen was approximately 3.3-fold higher in *chup1* KO plants compared to FNR control plants (Figures 1A and 1B), indicating that *chup1* KO plants are significantly more susceptible than the FNR control plants. Additionally, we generated independent *chup1* knockout lines from wild-type (WT) *N. benthamiana* plants (Figure S1F) (referred to as *chup1* KO#2 plants). These *chup1* KO#2 plants also demonstrated significantly higher *P. infestans* hyphal growth compared to the WT control plants (Figures 1C and 1D). These results implicate that the chloroplast outer envelope protein CHUP1 contributes to plant defense against adapted pathogens.

**Figure 1.**
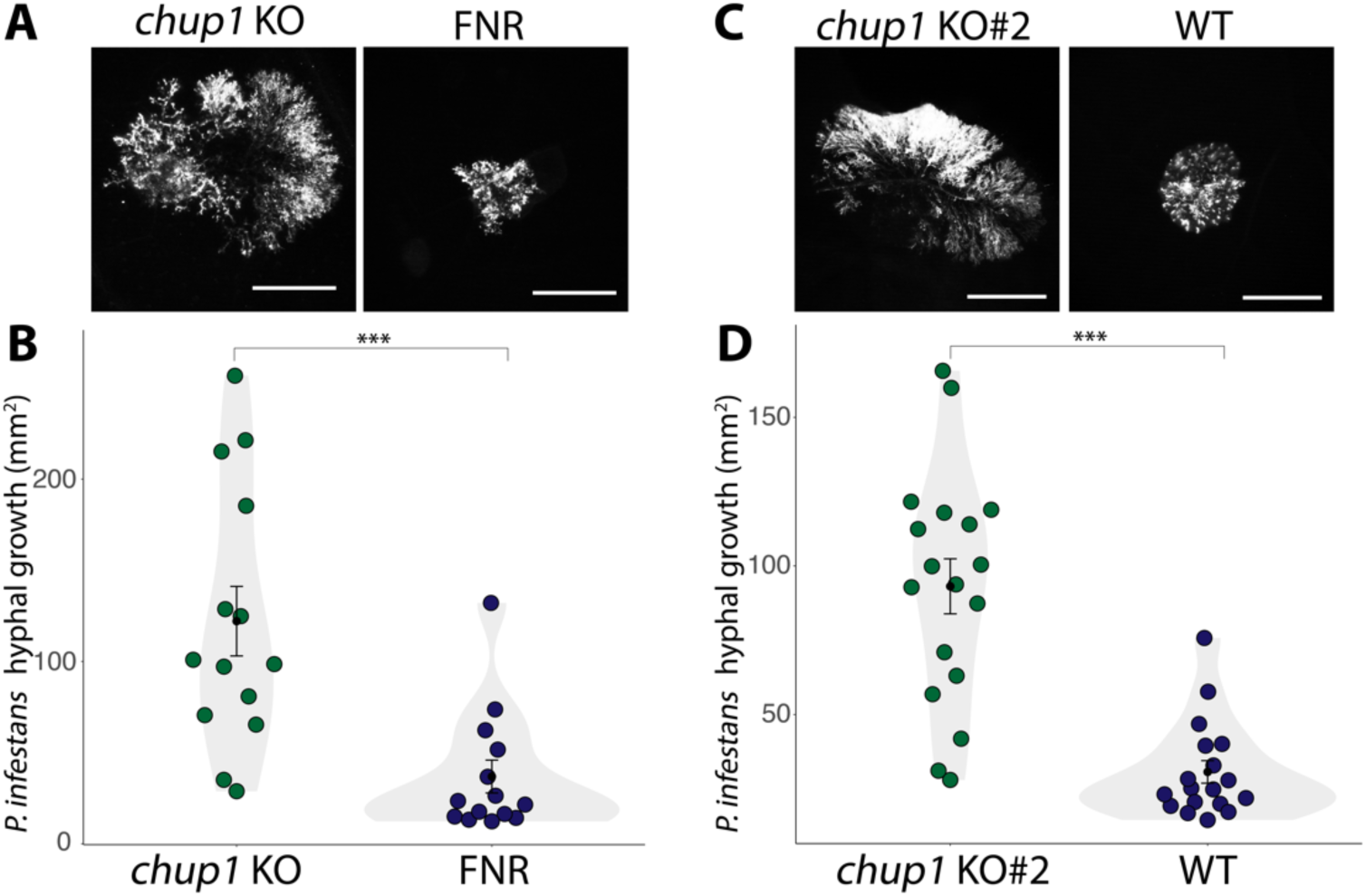
CHUP1 positively contributes to immunity against *P. infestans*. (A-B) *chup1* KO plants increase hyphal growth of *P. infestans* compared to control FNR plants. (A) 4-week-old *chup1* KO and FNR leaves were infected with tdTomato-expressing *P. infestans*, and pathogen growth was calculated by measuring hyphal growth using fluorescence stereomicroscope at 5 days post-inoculation. Scale bars represent 5 mm. (B) Violin plot illustrating that *chup1* KO plants (122.1 mm^2^, *n* = 83 infection spots) display a significant increase in *P. infestans* hyphal growth compared to control FNR plants (36.9 mm^2^, *n* = 83 infection spots). (C-D) *chup1* KO#2 plants increase hyphal growth of *P. infestans* compared to control WT plants. (C) 4-week-old *chup1* KO#2 and WT leaves were infected with tdTomato-expressing *P. infestans*, and pathogen growth was calculated by measuring hyphal growth using fluorescence stereomicroscope at 5 days post-inoculation. Scale bars represent 5 mm. (D) Violin plot illustrating that *chup1* KO#2 plants (93.1 mm^2^, *n* = 97 infection spots) exhibit a significant increase in *P. infestans* hyphal growth compared to control FNR plants (30.7 mm^2^, *n* = 94 infection spots). Each dot represents the average of all infection spots on the same leaf, with each leaf having between 3 to 6 infection spots. Statistical differences were analyzed by Mann-Whitney U test in R. Differences in measurements were considered highly significant when p<0.001 (***).

CHUP1 is known to facilitate chloroplast movement and photorelocation through cp-actin polymerization^31,37,38^. Therefore, we hypothesized that increased susceptibility in *chup1* KO plants could be due to perturbations in chloroplast movement and positioning around the haustoria. To investigate this, we imaged infected *N. benthamiana* epidermal cells and quantified chloroplast-haustoria associations in *chup1* KO and FNR plants. Our quantitative analysis showed no significant difference in the number of haustoria that is associated with chloroplasts between *chup1* KO and FNR plants (Figures S1G and S1H), indicating that enhanced susceptibility upon loss of CHUP1 is not due to impaired chloroplast movement towards the haustoria.

Chloroplasts clustering around nucleus is regarded to be a general plant immune response upon pathogen recognition and immune activation^18,20^. Therefore, we next checked whether chloroplast accumulation around the haustoria is impaired in *chup1* KO plants. Intriguingly, microscopy analysis revealed that chloroplasts tend to accumulate more around the nucleus in *chup1* KO plants compared to FNR plants (Figures S1I and S1J), indicating that the enhanced susceptibility of *chup1* KO plants is not due to impaired perinuclear clustering of chloroplasts. Lastly, we revealed that the number of chloroplasts in the epidermal cells and the total chlorophyll concentration in leaves did not vary significantly between *chup1* KO and FNR plants (Figures S1K-M). Altogether, these results indicate that CHUP1 is not essential for chloroplast positioning around haustoria, and the increased disease susceptibility in CHUP1 knockouts is not due to impaired chloroplast positioning or abnormal chloroplast numbers.

### CHUP1 accumulates at the chloroplast-plasma membrane and chloroplast-extrahaustorial membrane contact sites

To build on our findings of enhanced susceptibility in *chup1* KO plants, we next explored the cell biology of CHUP1 to better understand its role. We generated a C-terminal green fluorescent protein (GFP) fusion of CHUP1 (CHUP1:GFP) and examined its cellular localization using confocal microscopy. By co-expressing CHUP1:GFP with the PM marker REMORIN1.3 (REM1.3) tagged with red fluorescent protein (RFP), we observed that CHUP1:GFP accumulates at the chloroplast-PM membrane contact sites (MCS), forming puncta at this interface (Figure 2A). To validate this observation, we used Toc64:GFP as a control since both CHUP1 and Toc64 are chloroplast outer membrane proteins^39^. Unlike CHUP1:GFP, Toc64:GFP uniformly labeled the chloroplast outer membrane without forming punctate accumulations at chloroplast-PM MCS (Figure 2B). This contrast highlights the specific punctate localization pattern of CHUP1 at chloroplast-PM MCS.

**Figure 2.**
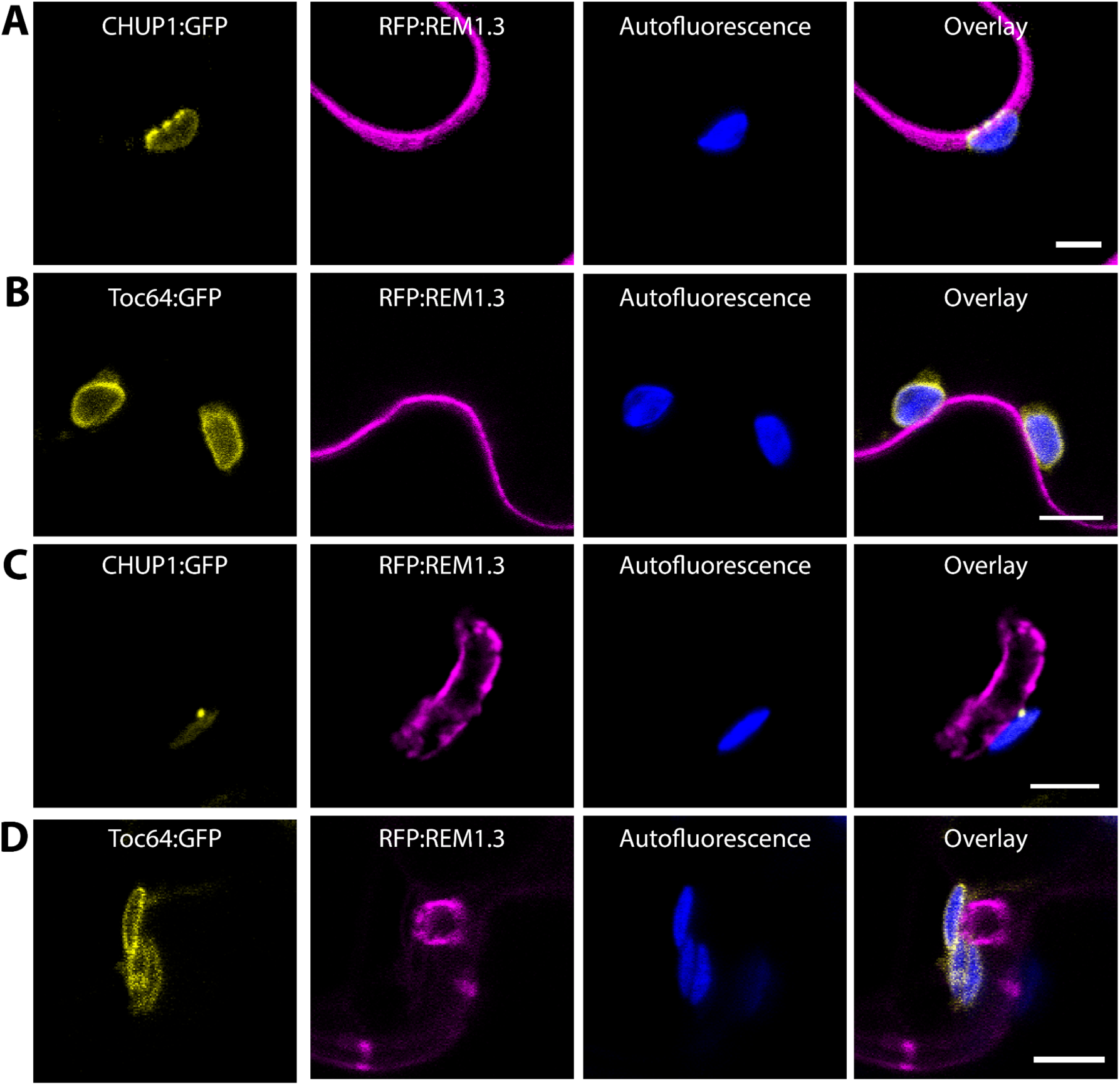
CHUP1 forms punctate accumulation at chloroplast-PM and chloroplast-EHM membrane contact sites (MCS). (A-B) CHUP1 forms punctate accumulation at chloroplast-PM MCS, but the control Toc64 does not. Confocal micrographs of *N. benthamiana* leaf epidermal cells transiently expressing either (A) CHUP1:GFP, or (B) Toc64:GFP, with RFP:REM1.3. Images were taken at 3 dpi. (C-D) CHUP1 forms punctate accumulation at chloroplast-EHM MCS, but the control Toc64 does not. Confocal micrographs of *N. benthamiana* leaf epidermal cells transiently expressing either (C) CHUP1:GFP, or (D) Toc64:GFP, with RFP:REM1.3. The leaves were infected with WT *P. infestans* spores at 6 hpi. Images were taken at 3 dpi. REM1.3 is used as a PM and EHM marker. Presented images are single plane images. Scale bars represent 5 µm.

Compared to the PM, the EHM has different lipid and protein contents, with most typical membrane proteins excluded from it^5,40,41^. Therefore, we next examined the localization of CHUP1, focusing on potential chloroplast-extrahaustorial membrane (EHM) MCS, in the context of *P. infestans* infection. Our live cell imaging of infected plant cells revealed that CHUP1:GFP also accumulates at foci where chloroplasts make contacts with the EHM (Figures 2C and S2A). In contrast, Toc64:GFP, used as a control, continued to show uniform labeling of the chloroplast outer membrane without any punctate accumulations at chloroplast-EHM MCS (Figures 2D and S2B). These observations reveal the distinctive accumulation of CHUP1 at MCS, both at chloroplast-PM and chloroplast-EHM, pointing to a role in chloroplast attachment to the pathogen interface.

### CHUP1 contributes to callose deposition at haustorium penetration sites

Recognizing that the increased disease susceptibility in *chup1* KO plants is not due to impaired chloroplast positioning around haustoria or abnormal chloroplast numbers (Figures S1G, S1H, S1K-M), and given that CHUP1 appears to anchor chloroplasts to the EHM (Figure 2C), we investigated whether focal plant immune responses towards the pathogen are altered in *chup1* KO plants. Accumulation of beta-glucan callose at haustoria penetration sites is a well-established focal immune response typically observed during plant invasion by fungal and oomycete pathogens^3,5,41^. To determine if callose deposition at the haustorium neck is affected in the absence of CHUP1, we employed aniline blue staining to visualize callose deposition around *P. infestans* haustorium in *chup1* KO and FNR plants (Figure 3A). Consistent with previous findings on accumulation of callose at the neckband of *P. infestans* haustoria^42^, approximately 20.4% of haustoria in infected FNR plants exhibited focal callose deposits (Figure 3B). Notably, we observed a 47.1% reduction in the number of haustoria with a callose neckband band in *chup1* KO plants, with only 10.8% of haustoria showing callose staining (Figure 3B). These results indicate that CHUP1 is implicated in focal callose deposition at the haustoria of *P. infestans*.

**Figure 3.**
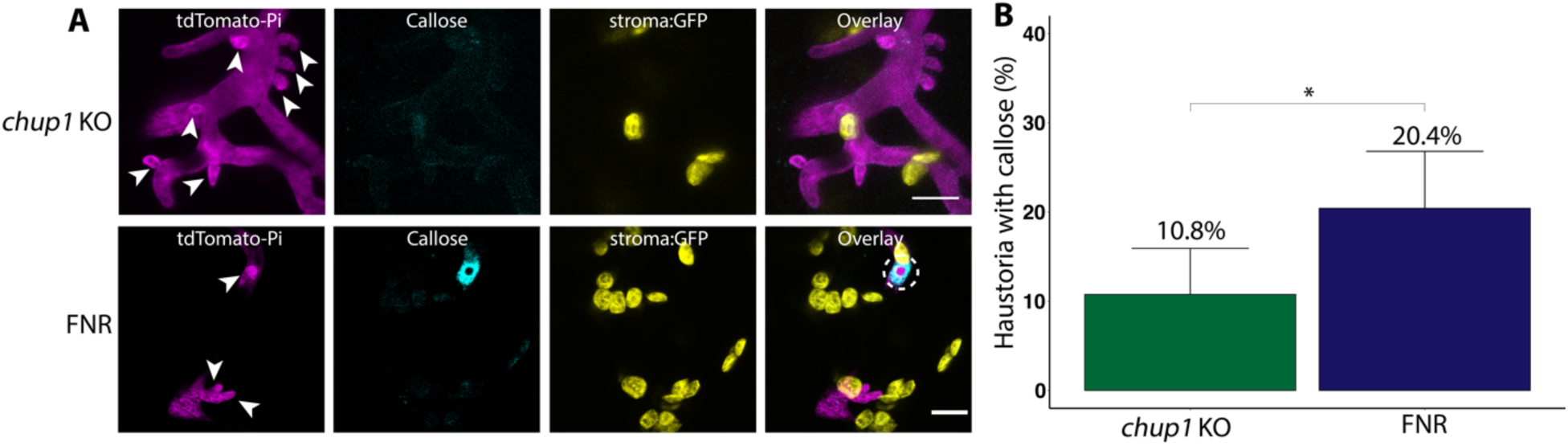
CHUP1 contributes to callose deposition at haustorium penetration sites. (A) Confocal micrographs of *N. benthamiana* leaf epidermal cells of *chup1* KO plants and control FNR plants. 4-week-old leaves were infected with tdTomato-expressing *P. infestans*, and stained with aniline blue to visualize callose at 3 days post-infection. Images shown are maximum projection of z-stack images. GFP channel depicts chloroplast stroma in *chup1* KO and FNR plants. White arrows indicate haustoria. Scale bars represent 10 μm. (B) Bar graphs showing *chup1* KO plants (10.8%, n = 130 haustoria) significantly reduce the frequency of callose deposition around haustoria compared to control FNR plants (20.4%, N = 142 haustoria). Statistical differences were analyzed by Fisher’s exact test in R. Differences in measurements were considered significant when p<0.05 (*).

We then investigated whether other core immune pathways were disrupted in *chup1* KO plants. To determine if basal immune responses following pathogen-associated molecular pattern (PAMP) recognition are altered in the absence of CHUP1, we measured mitogen-activated protein kinase (MAPK) phosphorylation after infiltrating leaves with *Phytophthora infestans* (Pi) extract^3^, which serves as a PAMP. Both *chup1* KO and FNR control plants showed a comparable increase in MAPK phosphorylation 24 hours post-PAMP treatment (Figure S3A). Next, we examined whether late-stage immune responses, such as defense-related cell death activation, were impaired in *chup1* KO plants. Following the expression of an autoactive variant of a MEK2-like NbMAPKK (MEK2^DD^)^43^, both *chup1* KO and FNR plants displayed a full tissue necrosis phenotype compared to the EV control (Figures S3B and S3C). We also tested whether effector-triggered immunity was impaired in the absence of CHUP1. To assess this, we co-expressed three cell death elicitors with their cognate NLR receptors in plants: *P. infestans* AVR3a with receptor R3a^44^, *P. infestans* effector AVRblb2 with receptor Rpi-blb2^45^, and potato virus X coat protein (PVX-CP) with receptor Rx^46^. Our cell death analysis showed that both *chup1* KO and FNR plants induced cell death to similar degrees in responses to AVR3a and R3a (Figures S3D and S3E), AVRblb2 and Rpi-blb2 (Figures S3F and S3G), and PVX-CP and Rx (Figures S3H and S3I). Altogether, these findings demonstrate that the activation of PAMP- or effector-triggered immunity is not impaired in the absence of CHUP1. We conclude that the enhanced susceptibility of *chup1* KO plants is most likely due to perturbations of MCS between chloroplasts and the EHM, leading to impaired focal immune responses as indicated by the reduced focal deployment of callose at the haustorium interface.

### KACs contribute to plant immunity against *P. infestans* and callose deposition around the haustoria

The kinesin-like proteins KACs have genetically overlapping and independent functions with CHUP1 in regulating chloroplast photorelocation^33,47^. Along with CHUP1, KACs are indispensable for the polymerization and maintenance of cp-actin filaments^33,34,48^. Through cp-actin filaments, these proteins are essential for both the proper movement of chloroplasts and their association with the PM^33^. While they coordinately mediate cp-actin-mediated chloroplast positioning, the underlying mechanism remains to be elucidated^38,47^.

The *N. benthamiana* genome includes two KAC genes, *KAC1* and *KAC2*. Given that CHUP1 and KACs have overlapping functions, we were intrigued to investigate if silencing KACs could confer a susceptibility phenotype similar to that observed when CHUP1 is knocked out. To explore this, we conducted infection assays upon downregulation of KAC expression. We employed a hairpin silencing construct, RNAi:KAC, designed to specifically target *KAC1* and *KAC2* in *N. benthamiana* (Figure S4). Similar to *CHUP1* silencing (Figures S1B and S1C), the silencing of *KACs* using RNAi:KAC significantly increased *P. infestans* hyphal growth compared to the control construct RNAi:GUS (Figures 4A and 4B). These results demonstrate that silencing *KACs* confers plant susceptibility, highlighting their role in plant immunity against *P. infestans*.

**Figure 4.**
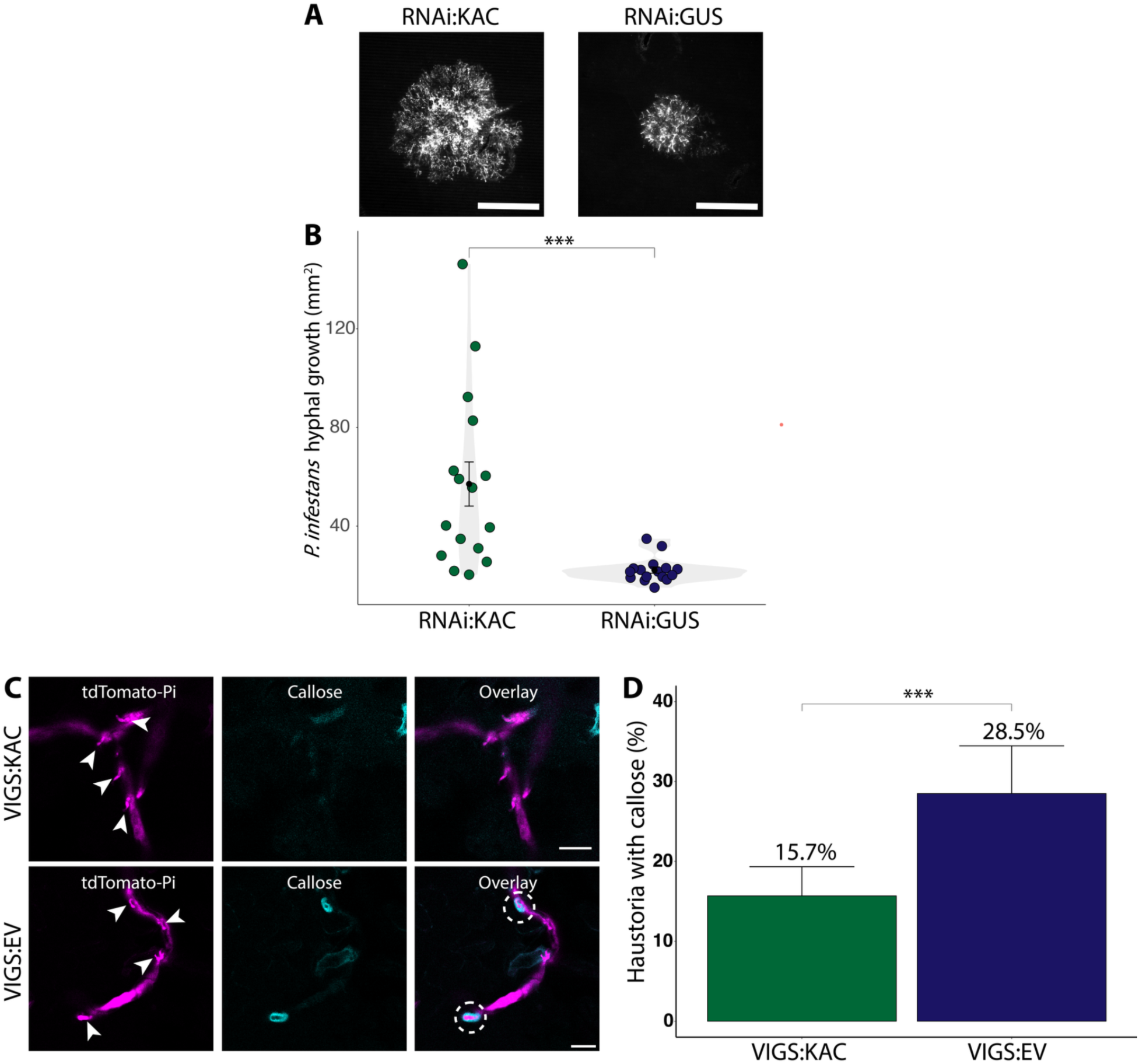
KACs positively contribute to immunity against *Phytophthora infestans* and reduce callose deposition surrounding *P. infestans* haustoria. (A-B) Silencing *KAC* reduces hyphal growth of *P. infestans*. (A) *N. benthamiana* leaves expressing RNAi:KAC, or RNAi:GUS control were infected with tdTomato-expressing *P. infestans*, and pathogen growth was calculated by measuring hyphal growth using fluorescence stereomicroscope at 5 days post-inoculation. Scale bars represent 5 mm. (B) Violin plot illustrating that RNAi:KAC expression (57.1 mm^2^, n = 48 infection spots) significantly increases *P. infestans* hyphal growth compared to RNAi:GUS control (22.1 mm^2^, n = 48 infection spots). Each dot represents the average of 3 infection spots on the same leaf. Statistical differences were analyzed by Mann-Whitney U test in R. Measurements were highly significant when p < 0.001 (***). (C-D) KAC proteins are involved in callose deposition at haustorium penetration sites. (C) Confocal micrographs of VIGS:KAC and VIGS:EV control *N. benthamiana* leaf epidermal cells. 4-week-old leaves were infected with tdTomato-expressing *P. infestans*, and stained with aniline blue to visualize callose at 3 days post-infection. Images shown are maximum projection of z-stack images. White arrows indicate haustoria. Scale bars represent 10 μm. (B) Bar graphs showing VIGS:KAC plants (15.69%, n = 376 haustoria) significantly reduce the frequency of callose deposition around haustoria compared to control VIGS:EV plants (28.51%, N = 221 haustoria). Statistical differences were analyzed by Fisher’s exact test in R. Differences in measurements were considered highly significant when p<0.001 (***).

We next investigated if KACs are also involved in focal immunity like CHUP1 (Figure 3). To accomplish this, we performed aniline blue staining of infected plant cells upon VIGS of KACs (Figure S4). We observed a reduction of approximately 44.9% (percentage change of frequency from 28.5% to 12.8%) in the number of haustoria with callose deposits in VIGS:KAC plants compared to VIGS:EV control plants (Figures 4C and 4D). These findings suggest that both CHUP1 and KAC proteins play a role in focal immunity and are involved in proper callose deposition at *P. infestans* haustoria.

### CHUP1 and KAC1 co-operate to tether chloroplasts to the pathogen interface

Having identified that both CHUP1 and KAC proteins play similar positive roles in immunity and are implicated in the focal immune responses against *P. infestans* haustoria, we aimed to explore their potential interplay in membrane tethering of chloroplasts. To address this, we first expressed C-terminally RFP-tagged KAC1 in *N. benthamiana* and investigated its localization pattern using confocal microscopy. Unlike CHUP1, which localizes to the chloroplast outer envelope (Figures 2A, 2C and S2A), KAC1:RFP mainly localizes to the PM and cytosol (Figure S5A). However, KAC1, similar to CHUP1 but not the control EV:RFP, accumulated at MCS between the chloroplasts and the PM (Figures S5B and S5C). Remarkably, in *chup1* KO plants, KAC1 did not show any accumulation at the chloroplast-PM contact sites (Figure S5D). This phenotype was restored by the complementation of CHUP1:3xHA co-expression, but not the co-expression of the empty vector control (3xHA:EV) (Figures S5D and S5E), demonstrating that KAC1 requires CHUP1 to accumulate at MCS between chloroplasts and the PM. These results indicate that KAC1 might have a direct role in membrane anchoring of chloroplasts in cooperation with CHUP1. This notion is consistent with the previous genetic studies implicating KAC1 and CHUP1 in chloroplast anchoring to the PM in a co-operative manner, as neither mutant was able to show anchorage independently^33,34^. Therefore, we next investigated the potential association between CHUP1 and KAC1. Co-expression of KAC1:RFP and CHUP1:GFP in *N. benthamiana* revealed that both proteins colocalize in punctate structures at MCS between chloroplasts and the PM (Figure S5F). In contrast, the controls, Toc64:GFP and EV:RFP, did not form any punctate structures at these MCS (Figures S5G and S5H), showing that not all chloroplast envelope proteins accumulate at the PM contact sites and indicating that CHUP1 and KAC1 cooperate to anchor chloroplasts to membranes.

Encouraged by these results, we next investigated KAC1-CHUP1 interplay in infected cells. In haustoriated plant cells, we observed that CHUP1 colocalizes with KAC1 at punctate structures at chloroplast-EHM MCS, whereas the empty vector control (EV:RFP) did not show such punctate localization (Figures 5A, 5B, S6A and S6B). Furthermore, upon infecting *chup1* KO plants expressing KAC1 with *P. infestans*, we found that KAC1 did not form puncta at chloroplast-EHM MCS (Figures 5C and S6C). Complementing *chup1* KO plants with CHUP1:3xHA reinstated KAC1 punctate formation at chloroplast-EHM MCS (Figures 5D and S6D). These findings collectively suggest that KAC1 requires CHUP1 to anchor chloroplasts to the plant-pathogen interface, presumably to execute their immune functions effectively. These results are consistent with our previous findings, which demonstrated that chloroplasts are tightly tethered to the EHM, as revealed by optical tweezer assays^7^.

**Figure 5.**
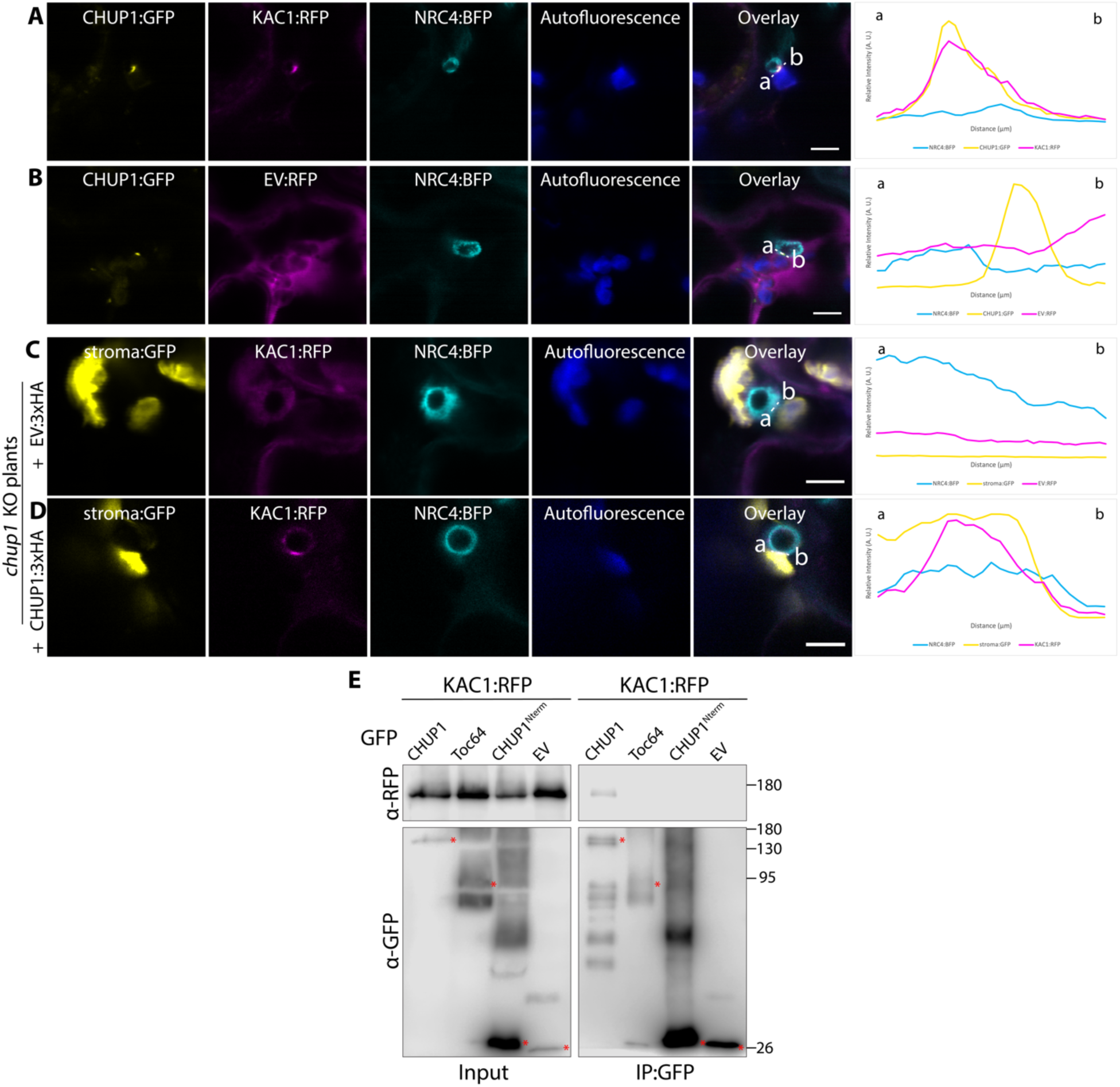
KAC1 interacts with CHUP1 and they colocalize at chloroplast-EHM MCS. (A-B) CHUP1 colocalizes with KAC1 at punctate structures at chloroplast-EHM MCS. Confocal micrographs of *N. benthamiana* leaf epidermal cells transiently expressing CHUP1:GFP and NRC4:BFP, with (A) KAC1:RFP, or (B) EV:RFP. NRC4:BFP acts as an EHM marker. The leaves were infected with WT *P. infestans* spores at 6 hpi, and imaged at 3 dpi. (C-D) CHUP1 is required for KAC1 to form punctate structures at chloroplast-EHM MCS. Confocal micrographs of *chup1* KO *N. benthamiana* leaf epidermal cells transiently expressing KAC1:RFP and NRC4:BFP with (C) EV:3xHA, or (D) CHUP1:3xHA. NRC4:BFP acts as an EHM marker. The leaves were infected with WT *P. infestans* spores at 6 hpi, and imaged at 3 dpi. All presented confocal images are single plane images. Autofluorescence channel depicts chloroplasts. GFP channel depicts chloroplast stroma in *chup1* KO plants. Transects in overlay panels correspond to line intensity plots depicting the relative fluorescence across the marked distance. Scale bars represent 5 µm. (E) KAC1 interacts with full length CHUP1, but not with Toc64, CHUP1^Nterm^, or EV:GFP. KAC1:RFP was transiently co-expressed with either CHUP1:GFP, Toc64:GFP, CHUP1^Nterm^:GFP, or EV:GFP. IPs were obtained with anti-GFP antibody. Total protein extracts were immunoblotted. Red asterisks indicate band sizes. Numbers on the right indicate kDa values.

To further corroborate the interplay between CHUP1 and KAC1, we next investigated their *in planta* interaction by performing co-immunoprecipitation (co-IP) assays. We co-expressed KAC1:RFP alongside CHUP1:GFP, or controls Toc64:GFP and CHUP1^Nterm^:GFP, the latter containing the N-terminal region of CHUP1 required for localization to chloroplast outer envelope^32^. Western blot analysis following GFP pulldowns revealed that KAC1:RFP interacts with CHUP1:GFP but not with Toc64:GFP, CHUP1^Nterm^:GFP, or EV:GFP (Figure 5E), corroborating our confocal microscopy findings that the two proteins associate at chloroplast MCS (Figures 5A, S5F and S6A).

CHUP1 is a modular protein that contains an actin-binding domain (ABD) necessary for the formation of chloroplast-actin filaments (cp-actin), short actin filaments that accumulate at the periphery of chloroplasts and the PM, which are implicated in chloroplast movement and PM anchoring^31,35^. Therefore, we lastly investigated whether the actin-binding function of CHUP1 is necessary for its complex formation with KAC1 and the formation of chloroplast-PM MCS. To address this, we generated GFP fusions of CHUP1ΔABD, a CHUP1 construct lacking the actin-binding domain. Confocal microscopy and co-IP assays revealed that the colocalization and interaction of CHUP1 and KAC1 at MCS do not depend on the actin-binding ability of CHUP1 (Figures S7A and S7B). However, we noted a slight reduction in the amount of CHUP1 pulled down with KAC1 when the actin-binding domain of CHUP1 was omitted (Figure S7B). We conclude that while the ABD of CHUP1 is not essential for CHUP1-KAC1 interaction and their assembly at chloroplast MCS, it could be important for coupling them more tightly to ensure secure docking of chloroplasts at the PM and EHM. Collectively, our findings highlight that KAC1 interacts with CHUP1 and both proteins colocalize at crucial membrane contact sites, facilitating the anchoring of chloroplasts to the PM and the plant-pathogen interface.

## Discussion

Here, we uncovered MCS between chloroplasts and the EHM in *P. infestans*-infected plant cells. We reveal that these MCS are required for proper immune responses and consist of a membrane anchoring complex comprising the chloroplast outer envelope protein CHUP1 and the PM-associated protein KAC1. Our genetic and cell biology analyses revealed that KAC1 marks these MCS only in the presence of CHUP1. Additionally, our biochemical assays show that CHUP1 and KAC1 interact. We propose a model in which CHUP1 and KAC1 synergistically contribute to plant focal immunity by facilitating MCS at the pathogen penetration sites during infection (Figure 6).

**Figure 6.**
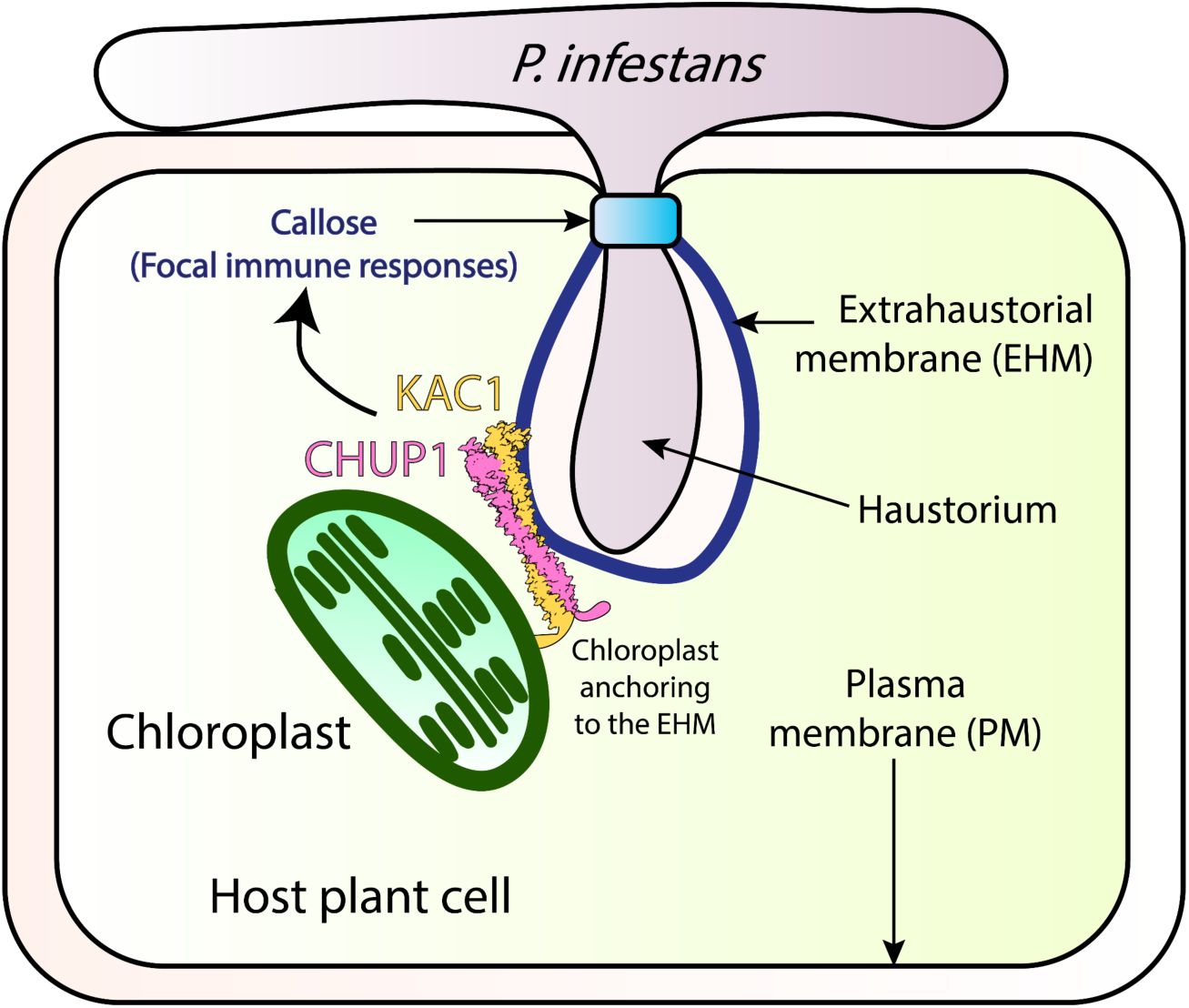
Summary model of CHUP1 and KAC1 coordinating to anchor chloroplasts to the extrahaustorial membrane and contribute to plant focal immunity. *P. infestans* penetrates the host plant cell by forming a specialized structure called the haustorium, which is surrounded by a host-derived extrahaustorial membrane (EHM) that acts as the plant-pathogen interface. The chloroplast movement-associated proteins CHUP1 and KAC1 interact to form chloroplast-EHM membrane contact sites (MCS), thereby anchoring chloroplasts to the EHM. This interaction contributes to plant focal immunity, such as callose deposition and potentially other immune responses at the plant-pathogen interface.

The role of CHUP1-KAC1 MCS in pathogen resistance is underscored by our findings that depletion of CHUP1 or KAC1 leads to enhanced susceptibility to the adapted pathogen *P. infestans* (Figures 1 and 4). This susceptibility is accompanied by a reduction in focal immune responses, as evidenced by decreased callose accumulation around the haustorium, without affecting other core immune processes (Figures 3, 4 and S3). These results align with findings from other studies showing that *CHUP1* knockouts in Arabidopsis similarly result in decreased penetration resistance to non-adapted pathogens^36^. Additionally, previous studies have implicated chloroplast outer envelope proteins AtNHRA/B in the proper accumulation of callose^49^, further supporting our observations that chloroplasts contribute to callose accumulation around the haustorium.

Prior research has also revealed that mitochondria accumulates at pathogen penetration sites and contribute to pathogen penetration resistance, although it remains unknown whether mitochondria can associate with the EHM^8^. Our previous work using optical tweezers showed that chloroplasts can be tightly tethered to the EHM^7^, but the mechanisms and reasons for this remained unknown. While the precise mechanism by which CHUP1-KAC1 mediates callose deposition around the haustorium remains unknown, their role in tethering chloroplasts to the EHM suggests a significant involvement of chloroplasts in resistance to pathogen penetration. These tethers could ensure secure membrane contacts, facilitating potential short distance material transport through chloroplast-EHM conduits. Indeed, one of the key features of MCS is the direct transfer of lipids, proteins, signaling molecules, ions and metabolites. It is conceivable that CHUP1-KAC1-labeled MCS facilitate the transport of biochemical precursors necessary for callose synthesis or the transfer of ROS that cross-link and fortify callose deposits^50–52^. These MCS may act as critical hubs for the exchange of materials and signals essential for reinforcing the cell wall at pathogen entry points.

Our current understanding of the mechanisms of chloroplast movement and membrane anchoring originates from studies focusing on chloroplast photorelocation, a process for maximizing photosynthesis and minimizing photodamage based on exposure to light intensities. While dozens of proteins are implicated in chloroplast movement, the detailed molecular interactions and mechanisms remain largely elusive. Previous genetic studies have identified CHUP1 and KAC1 as key players in chloroplast photorelocation^33,47,53^. Our findings align with this, showing that CHUP1 accumulates at foci where chloroplasts contact the PM (Figure 2A). Additionally, we observed that CHUP1 also localizes at EHM contact sites (Figure 2C), and that KAC1 accumulates at these sites in a CHUP1-dependent manner (Figures 5C, 5D, S5D and S5E). Our biochemical assays further support the physical interaction between CHUP1 and KAC1 (Figures 5E and S7B). These results suggest that the machinery involved in chloroplast photorelocation is co-opted for innate immune responses. Furthermore, blue light receptors, which regulate chloroplast photorelocation, are also implicated in plant immunity^54,55^, underscoring the multifaceted role of these components in both light response and pathogen defense mechanisms.

The exact biochemical and molecular mechanisms through which CHUP1 and KAC1 facilitate MCS tethering chloroplasts to the pathogen interface and contribute to pathogen resistance are yet to be fully elucidated. Future studies are required to determine how these MCS formed by CHUP1 and KAC1 enhance immune responses. Investigating the dynamics of these contact sites and their precise molecular and biochemical functions will be crucial in understanding how they contribute to the targeted deposition of defense components at the pathogen interface. Understanding the intricacies of MCS at the pathogen interface will provide deeper insights into plant defense mechanisms and could inform the development of new strategies to enhance crop disease resistance.

## Materials and Methods

### Plant growth details

*Nicotiana benthamiana* plants (wild-type and transgenics) were cultivated under controlled conditions in a growth chamber maintained at a temperature of 24°C. They were grown in a substrate mixture comprising organic soil (3:1 ratio of Levington’s F2 with sand and Sinclair’s 2-5 mm vermiculite). The plants were exposed to high-intensity light and subjected to a long-day photoperiod (16 hours of light followed by 8 hours of darkness). Experiments were conducted using plants aged 4 to 5 weeks. Among the transgenic plants utilized were CP plants^7^, expressing GFP in the chloroplast stroma, and FNR plants^56^, expressing FNR:eGFP in the chloroplast. FNR plants served as the parental lines for *chup1* KO plants and were employed as controls in *chup1* KO experiments.

### Pathogen growth details and infection assay

WT and tdTomato-expressing *Pytophthora infestans* 88069 isolates were cultured on rye sucrose agar (RSA) medium in darkness at 18°C for 10 to 15 days before harvesting zoospores. Zoospores were collected by adding cold water at 4°C to the medium and then incubating it in darkness at 4°C for 90 minutes. For the infection assay, 10 µl droplets of zoospore solution containing 50,000 spores/ml were applied to the underside of leaves. The leaves were subsequently maintained in a humid environment. Confocal microscopy was conducted 3 days post-infection, and fluorescent images were captured at 5 days post-infection. Hyphal growth was quantified and analyzed using ImageJ.

### Generation of *N. benthamiana* CHUP1 CRISPR knockout plants

To generate *chup1* KO and *chup1* KO#2 plants, the following primer pairs were used to create guide RNA sequences targeting both *NbChup1* alleles: attGCAAGATCAAGGAGTTGCAG & aaacCTGCAACTCCTTGATCTTG; attgTGGACTTCAAGAAAAGGAAG & aaacCTTCCTTTTCTTGAAGTCCA; and attgTCTGTATCATACTTGTCACT & aaacAGTGACAAGTATGATACAGA. The genome editing vector used was pDGE463^57^. This vector contains: plant kanamycin resistance for positive selection of transgenic plants, the Bs3 gene from pepper for negative selection, 2xtagRFP expressed by a seed coat-specific promoter for negative selection, a p35S-driven intronized Cas9 with 2xNLS for targeted gene editing, and bacterial spectinomycin resistance for selection in bacteria. The guide RNAs were inserted into pDGE463 to create the new vector, pDGE472. The primers used for genotyping *chup1* KO plants were CHUP1KO_genotype_F and CHUP1KO_genotype_R, while the primers used for genotyping *chup1* KO#2 plants were CHUP1KO2_genotype_F and CHUP1KO2_genotype_R. The parental line transformed was NbFNR:eGFP_7-25 (FNR) for *chup1* KO plants^58^, and the parental line transformed was wild-type (WT) for *chup1* KO#2 plants.

### Molecular cloning

The molecular cloning of KAC1, CHUP1, CHUP1^Nterm^, CHUP1ΔABD, MEK2^DD^, and NLS was conducted using Gibson Assembly, following the methods described in previous works^3,59^. Specifically, the vector backbone was a pK7WGF2 derivative domesticated for Gibson Assembly, containing a C-terminal fluorescent GFP, RFP, BFP, or 3xHA tag. The desired sequences for cloning were either manufactured as a synthetic fragment or amplified using designed primers as detailed in Table S1 and S2. CHUP1ΔABD was created using two fragments that were amplified with two sets of primers, excluding the actin-binding domain flanked by the fragments. The fragments were then inserted into the vector using Gibson Assembly and transformed into DH5α chemically competent *E. coli* through heat shock. These plasmids were subsequently amplified and extracted using the PureYield™ Plasmid Miniprep System (Promega), and electroporated into *Agrobacterium tumefaciens* GV3101 electrocompetent cells. Sequencing was done by Eurofins. LB agar containing gentamicin and spectinomycin was used to grow bacteria carrying the pK7WGF2 plasmid. The NLS:BFP construct was created by joining a cut pK7WGF2-derived C-terminal BFP vector with a single-stranded DNA oligo bridge using Gibson assembly. The single-stranded DNA oligo bridge had overhangs complementary to the cut vector ends and included the coding sequence for the SV40 nuclear localization sequence (PKKKRKVEDP). For virus-induced gene silencing (VIGS), TRV2:CHUP1 was constructed by amplifying a region of the CHUP1 sequence using the primer pair CHUP1_sil_F and CHUP1_sil_R. The amplified fragment was then cloned into a Gateway-compatible pTRV2 vector using Gateway cloning technology (Invitrogen). TRV2:KAC for VIGS was constructed by amplifying a region of the KAC1 sequence using the primer pair KAC1_sil_F and KAC1_sil_R. The amplified fragment was then cloned into a Golden Gate-compatible TRV2-GG vector using Golden Gate cloning^60^. LB agar containing gentamicin and kanamycin was used to grow bacteria carrying pTRV2 and TRV2-GG plasmids. For the RNA interference (RNAi) silencing construct RNAi:KAC, an intron-containing hairpin RNA vector for RNA interference in plants (pRNAi-GG) was employed, based on the Golden Gate cloning method described in previous studies^3,61^. RNAi:KAC targeted the region between 3512 and 3709 bp of *NbKAC1/2*. The target fragment was amplified using KAC1_sil_F and KAC1_sil_R and then inserted into the pRNAi-GG vector in both sense and antisense orientations, utilizing the overhangs left by BsaI cleavage. This resulted in the expression of a construct that folds back onto itself, forming the silencing hairpin structure. The subsequent steps of *E. coli* transformation, Miniprep, sequencing, and Agrobacterium transformation were the same as those used for the overexpression constructs. LB agar containing gentamicin, kanamycin, and chloramphenicol was used to grow bacteria carrying the pRNAi-GG plasmid. All primers and synthetic fragments used in this study are detailed in Table S1. All constructs used in this study are detailed in Table S2.

### Confocal laser scanning microscopy

The confocal microscopy analyses were conducted three days following agroinfiltration. To visualize the leaf tissue, the leaves were excised using a size 4 cork borer, mounted live on glass slides, and submerged in wells of dH_2_O using Carolina observation gel (Carolina Biological). Imaging of the abaxial side of the leaf tissue was performed using either a Leica TCS SP8 inverted confocal microscope equipped with a 40x water immersion objective lens or a Leica STELLARIS 5 inverted confocal microscope equipped with a 63x water immersion objective lens. Laser excitations for GFP, RFP/tdTomato, and BFP tags were set at Argon 488 nm (15%), DPSS 561 nm, and Diode 405 nm, respectively. Emission ranges for GFP, RFP/tdTomato, and BFP tags were 495 - 550 nm, 570 - 620 nm, and 402 - 457 nm, respectively. To prevent spectral overlap from different fluorescent tags when imaging samples with multiple tags, sequential scanning between lines was employed. Confocal images were analyzed using ImageJ.

### Confocal Image analysis

The number of chloroplasts was automatically counted using ImageJ. Maximum intensity z-projections of z-stack images were created, and the channels were split. The autofluorescence channel was then automatically thresholded using the “Li” method, followed by the application of a watershed. Objects were counted using the “Analyze Particles” command with a size filter of 5-35.

For quantifying chloroplast-haustoria association, haustoria were first identified in a given z-stack using only the RFP and brightfield channels to ensure that their identification was not influenced by the position of chloroplasts. The number of haustoria associated with a chloroplast, including those connected via a stromule, was then counted. The percentage of haustoria associated with chloroplasts was calculated based on the total number of identified haustoria. To avoid skewing the data with images containing a single haustorium, which could result in extreme percentages (0% or 100%), the overall percentage for the entire data set was used instead of calculating percentages on an image-by-image basis.

Quantifying perinuclear chloroplast clustering was challenging due to the close packing of chloroplasts around the nucleus, making it difficult to resolve individual chloroplasts. To measure perinuclear clustering, the total volume of thresholded chlorophyll autofluorescence around the nuclei was quantified. First, nuclei were automatically identified from maximum intensity z-projections of z-stack images, with channels split. The NLS:BFP channel was automatically thresholded using the “Li” method, followed by the application of a watershed. Nuclei were then counted using the “Analyze Particles” command with a size filter of 100-500. The positions of the identified nuclei were saved as regions of interest (ROIs). For each ROI, the original image was duplicated and cropped to a 70x70 pixel region around the nucleus, which was then saved separately for further analysis. The quality of these cropped images was manually assessed. If necessary, the following adjustments were made: a) Slices of the z-stack containing the mesophyll layer were removed. b) Crops were resized to exclude chloroplasts from adjacent cells or those not in immediate contact with the nucleus. The chloroplast autofluorescence channel was then isolated and automatically thresholded using the “Li” method. For each slice of the z-stack, pixels with a value of 255 were counted, and the voxel size was determined for each image. This data was compiled into a custom table containing pixel count, voxel size, and image name for all identified nuclei. Using this table, the cubic microns of thresholded chloroplast autofluorescence were calculated for each identified nucleus, providing a metric for perinuclear chloroplast clustering.

### Fluorescence microscopy for *N. benthamiana*

Fluorescence microscopy was employed to observe the hyphal growth of *P. infestans* expressing tdTomato. The imaging setup included a Leica MZ 16 F microscope paired with the Leica DFC300 FX Digital Color Camera, specifically designed for fluorescence imaging. Infected leaf samples were placed on a petri dish within the imaging zone of the microscope. The imaging filter utilized was DsRed, with an excitation range ranging from 510 to 560 nm.

### Agrobacterium-mediated transient gene expression in *N. benthamiana*

Agrobacterium-mediated transient gene expression was carried out via agroinfiltration, following a well-established protocol^1^. *Agrobacterium tumefaciens* containing the desired plasmid was rinsed with water and then suspended in agroinfiltration buffer (10 mM MES, 10 mM MgCl_2_, pH 5.7). The optical density (OD_600_) of the bacterial suspension was measured using the BioPhotometer spectrophotometer (Eppendorf). Subsequently, the suspension was adjusted to the required OD_600_ according to the specific construct and experimental conditions. The adjusted bacterial suspension was then infiltrated into *N. benthamiana* leaf tissue aged 3 to 4 weeks using a needleless 1ml Plastipak syringe.

### RNA isolation, cDNA synthesis, and RT-PCR

To perform RNA extraction, 56 - 70 mg of leaf tissue was promptly frozen in liquid nitrogen. The RNA extraction process employed the Absolutely Total RNA Purification Kits (Agilent Technologies). Subsequently, RNA concentration was quantified using NanoDrop Lite Spectrophotometer (Thermo Scientific). 2 mg of the extracted RNA underwent treatment with RQ1 RNase-Free DNAse (Promega) before being used for cDNA synthesis with SuperScript IV Reverse Transcriptase (Invitrogen). The resulting cDNA was then amplified using Phusion High-Fidelity DNA Polymerase (New England Biolabs) or DreamTaq DNA polymerase (Thermo Scientific). *GAPDH* transcription level was utilized as the transcriptional control. The RT-PCR for *GAPDH* was performed using the primers GAPDH_RTPCR_F and GAPDH_RTPCR_R. The RT-PCR confirming VIGS:CHUP1 was performed using the primers NbCHUP1a_RTPCR_F and NbCHUP1a_RTPCR_R for *NbCHUP1a* and NbCHUP1b_RTPCR_F and NbCHUP1b_RTPCR_R for *NbCHUP1b*. The RT-PCR confirming RNAi:KAC and VIGS:KAC was performed using the primers KAC1_RTPCR_F and KAC1_RTPCR_R for *NbKAC1* and KAC2_RTPCR_F and KAC2_RTPCR_R for *NbKAC2*. All primers used in this study are detailed in Table S1.

### Chlorophyll extraction

Three leaf discs, obtained with a size 4 cork borer, were collected from three distinct leaves of each plant. Each leaf disc was then ground in 10 mL of methanol for 1 minute and centrifuged at 2000 rpm for 5 minutes. Absorbance readings at 666 and 653 nm were recorded, and the total chlorophyll concentration was determined following a well-established method detailed by Wellburn et al^62^.

### Co-immunoprecipitation and immunoblot analyses

Proteins were transiently expressed in *N. benthamiana* leaves through agroinfiltration, and the harvest took place 3 days after agroinfiltration. Co-immunoprecipitation experiments utilized 2 g of leaf tissues. The procedures for protein extraction, purification, and immunoblot analysis followed the previously described protocols^1^. The primary antibodies used included monoclonal anti-RFP produced in mouse (Chromotek), monoclonal anti-GFP produced in rat (Chromotek), polyclonal anti-phospho-MAPK produced in rabbit (Cell Signaling Technology), and monoclonal HRP-conjugated anti-beta actin produced in mouse (Proteintech). As for secondary antibodies, anti-rabbit antibody (Sigma-Aldrich), anti-rat antibody (Sigma-Aldrich), and anti-mouse antibody (Sigma-Aldrich) for HRP detection were employed. Comprehensive information regarding the antibodies used is detailed below.

List of antibodies used in this research:

| Primary/Secondary | Detail | Company | Product Code |
| --- | --- | --- | --- |
| Primary | Anti-GFP (Rat monoclonal) | Chromotek | 3H9 |
| Primary | Anti-RFP (Mouse monoclonal) | Chromotek | 6G6 |
| Primary | Anti-phospho-MAPK (Rabbit polyclonal) | Cell Signaling Technology | 9101 |
| Secondary | Anti-rabbit (HRP) | Sigma-Aldrich | A9169 |
| Secondary | Anti-rat (HRP) | Sigma-Aldrich | A9037 |
| Secondary | Anti-mouse (HRP) | Sigma-Aldrich | SAB3700986 |
| Primary/Secondary | HRP-conjugated anti-beta actin<br>(Mouse monoclonal) | Proteintech | HRP-60008 |

### Cell death assay

Cell death inducers were introduced into the underside of *N. benthamiana* leaves via agroinfiltration. Afterward, at 3 dpi, the leaves were detached and examined under both natural light conditions. The degree of cell death was assessed using an established seven-tiered cell death scale^63^.

### *Phytophthora infestans* extract (Pi extract) preparation and injection

Mycelia harvested from *P. infestans* RSA plates were gathered and suspended in 5 ml of water per petri dish. The suspension underwent vortexing for 1 minute and was then subjected to heating at 95°C for 20 minutes. Following this, the mixture was filtered through filter paper with a pore size ranging from 5 to 13 µm. The resulting filtrate underwent an additional filtration step using a syringe filter with a pore size of 0.45 µm. This resulting solution was then administered to plants to function as a PAMP cocktail, and is stored at -20°C.

### Virus-induced gene silencing (VIGS)

Agrobacterium was prepared as described above, carrying TRV1 and the respective TRV2 construct, and mixed to achieve final OD_600_ values of 0.4 or 0.2, respectively, in agroinfiltration buffer supplemented with 100 µM acetosyringone (Sigma-Aldrich). The mixture was then kept in the dark for 2 hours prior to infiltration to enhance virulence. Fourteen-day-old *N. benthamiana* seedlings were infiltrated in both cotyledons and any emerged true leaves. For CHUP1-silencing, *N. benthamiana* plants were infiltrated with TRV1 and TRV2:CHUP1, while for KAC1-silencing, TRV1 and TRV2:KAC were used. The empty vector control was achieved by infiltrating TRV1 and TRV2:EV. Plants were allowed to grow under standard conditions until experiments could be conducted three weeks later.

### Image processing and data analysis

The confocal microscopy images were processed using Leica LAS X software and ImageJ. ImageJ was employed for analyzing and quantifying the infection assay experiments. Data were presented using violin plots, box plots and bar graphs created with R. Statistical differences were evaluated using appropriate tests that consider normality and variance. Significance levels were denoted as follows: * (p < 0.05), ** (p < 0.01), and *** (p < 0.001). Detailed information regarding the statistical tests utilized can be found in the figure captions, and extensive statistical calculations are available in Table S3.

### Accession numbers

CHUP1a (Sol Genomics Network: Niben101Scf00570g03008/9.1; NbenBase: Nbe.v1.1.chr08g25910); CHUP1b (Sol Genomics Network: Niben101Scf01338g03019.1); KAC1 (NbenBase: Nbe.v1.1.chr13g42800); KAC2 (NbenBase: Nbe.v1.1.chr05g36640).

## Supporting information

Summary of Statistics

## Acknowledgments

We gratefully acknowledge the technical expertise and access to microscopy equipment provided by the Imperial College FILM facility.

## Author contributions

Conceptualization: E.L.H.Y., Z.S., M.S., T.O.B.

Methodology: E.L.H.Y., Z.S., M.S., T.O.B.

Validation: E.L.H.Y., Z.S., V.A., C.V., Y.Z., F.R., A.I.B.

Formal Analysis: E.L.H.Y., Z.S., V.A., C.V., Y.Z., A.I.B.

Investigation: E.L.H.Y., Z.S., V.A., C.V., Y.Z., F.R., A.I.B.

Resources: E.L.H.Y., Z.S., V.A., Y.T., J.L.E., J.P., J.S., C.D., C.M.

Data Curation: E.L.H.Y., T.O.B.

Visualization: E.L.H.Y., Z.S.

Writing – Original Draft: E.L.H.Y., Z.S., T.O.B.

Writing – Review & Editing: E.L.H.Y., Y.T., T.O.B.

Supervision: E.L.H.Y., Z.S., M.S., T.O.B.

Funding Acquisition: M.S., T.O.B.

## Declaration of generative AI and AI-assisted technologies in the writing process

During the preparation of this work, the authors used ChatGPT to check grammar and phrasing. After using ChatGPT, the authors reviewed and edited the content as needed and take full responsibility for the content of the publication.

## Data and materials availability

All relevant study data are included in the article, and in the Supplementary Materials files.

## Supplementary Materials in this combined PDF include the following

Supplementary Figures S1-S7

Supplementary Tables S1-S2

## Other Supplementary Materials for this manuscript include the following

Table S3. Summary of Statistics

---

**Figure S1.**
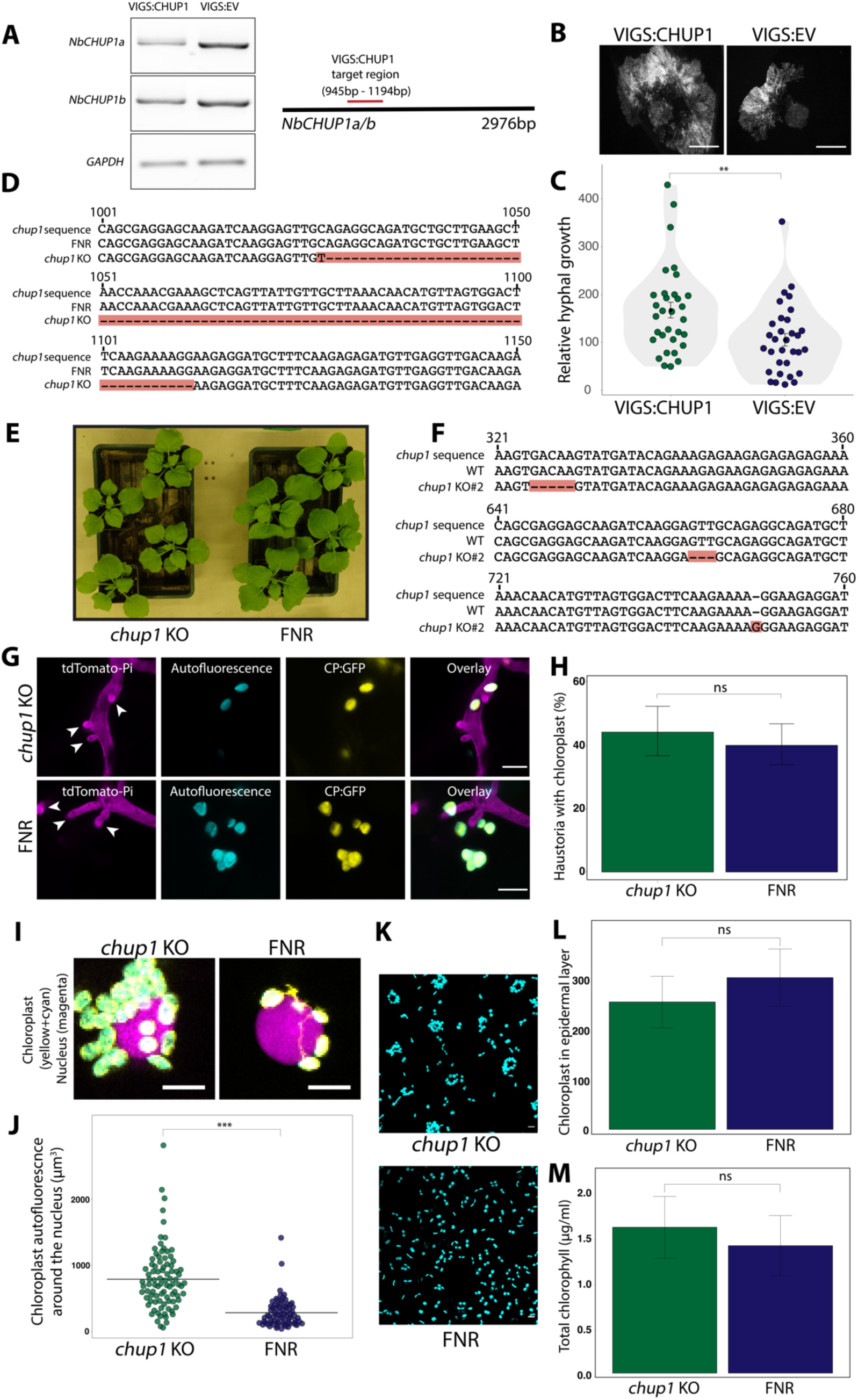
CHUP1 positively contributes to plant immunity and the increased disease susceptibility observed in *chup1* KO plants is not attributed to impaired chloroplast positioning or abnormal chloroplast numbers. (A) Validation of *CHUP1* silencing of VIGS:CHUP1 plants. Constructs carrying TRV1 with TRV2-GG targeting *NbCHUP1a and NbCHUP1b* or the control TRV2:EV were infiltrated to *N. benthamiana* leaves. The expression levels of the targeted genes were assessed via RT-PCR at 3 weeks post VIGS. The RT-PCR employs primers NbCHUP1a_RTPCR_F and NbCHUP1a_RTPCR_R for *NbCHUP1a*, and NbCHUP1b_RTPCR_F and NbCHUP1b_RTPCR_R for *NbCHUP1b*. RT-PCR results confirmed gene silencing of *NbCHUP1a* and *NbCHUP1b* in the VIGS:CHUP1 plants. Glyceraldehyde 3-phosphate dehydrogenase (GAPDH) served as the internal control, using primers GAPDH_RTPCR_F and GAPDH_RTPCR_R for assessment. The cDNA was synthesized using total RNA. (B) WT *N. benthamiana* leaves were infected with tdTomato-expressing *P. infestans* 3 weeks post VIGS, and pathogen growth was calculated by measuring hyphal growth using fluorescence stereomicroscope at 5 days post-inoculation. Scale bars represent 5 mm. (C) Violin plot illustrating that plants treated with VIGS:CHUP1 construct (167.7, *n* = 192 infection spots) show a significant increase in *P. infestans* hyphal growth compared to plants treated with control VIGS:EV construct (105.6, *n* = 192 infection spots). Each dot represents the average of 6 infection spots on the same leaf. Statistical differences were analyzed by Mann-Whitney U test in R. Measurements were significant when p<0.01. (**). (D) Generation of *N. benthamiana chup1* CRISPR knockout mutant, designated as *chup1* KO plants, in an FNR:eGFP background. *chup1* KO plants contain a 84 nt deletion that introduces premature stop codons in the *chup1* gene. (E) Daylight photographs of representative four-week-old *chup1* KO and FNR plants used for experiments. (F) Generation of *N. benthamiana chup1* CRISPR knockout mutant, designated as *chup1* KO#2 plants, in a WT background. *chup1* KO#2 plants contain three editing sites in the *chup1* gene. First site contains a 5 nt deletion and causes a frameshift with stop codons downstream. Second site contains a 3 nt deletion. Third site contains a 1 nt insertion. These mutations do not reconstitute the reading frame. (G-H) Chloroplast positioning at haustoria is unaffected by the loss of CHUP1. (G) Confocal micrographs depict *chup1* KO and FNR control *N. benthamiana* leaf epidermal cells. Four-week-old leaves were infected with tdTomato-expressing *P. infestans*. Imaging was performed 3 days post-inoculation. White arrows indicate haustoria. Yellow and cyan show chloroplast stroma and chloroplast autofluorescence respectively. Scale bars represent 10 µm. (H) Bar graphs demonstrate that the differences in chloroplast-haustoria associations between *chup1* KO plants (44.38%, *n* = 160 haustoria) and FNR plants (40.17%, *n* = 229 haustoria) are not significant. Error bars represent standard deviation. Statistical significance was assessed using Fisher’s exact test in R. (I-J) Chloroplasts in *chup1* KO plants are clustered around the nucleus. (I) Confocal micrographs depicting nuclei in *chup1* KO and FNR plants transiently expressing NLS:BFP (magenta). Yellow and cyan show chloroplast stroma and chloroplast autofluorescence respectively. Scale bars represent 10 µm. (J) Quantification of chloroplast autofluorescence volume surrounding nuclei, marked by transiently expressed NLS:BFP, in *chup1* KO and FNR plants. *chup1* KO plants (791, *n* = 89 nuclei) exhibit a significant increase in chloroplast clustering around the nucleus compared to control FNR plants (286, *n* = 74 nuclei). Crossbars represent the mean. Statistical significance was determined using the Mann-Whitney U test in R. Each data point represents a measurement from a single isolated nucleus. (K) Confocal micrographs depict *chup1* KO and FNR plants where the entire depth of the epidermal layer is shown. Chloroplasts are depicted in cyan colour. (L) Bar graphs demonstrate that the differences in the number of chloroplasts in the epidermal layer between *chup1* KO plants (253, *n* = 9 micrographs) and FNR plants (301, *n* = 0 micrographs) are not significant. Error bars represent standard deviation. Statistical significance was assessed using Student’s t-test in R. (M) Bar graphs demonstrate that the differences in the total chlorophyll concentration between *chup1* KO plants (1.6 µg/ml, *n* = 3 plants) and FNR plants (1.4 µg/ml, *n* = 3 plants) are not significant. Error bars represent standard deviation. Statistical significance was assessed using Mann-Whitney U test in R.

**Figure S2.**
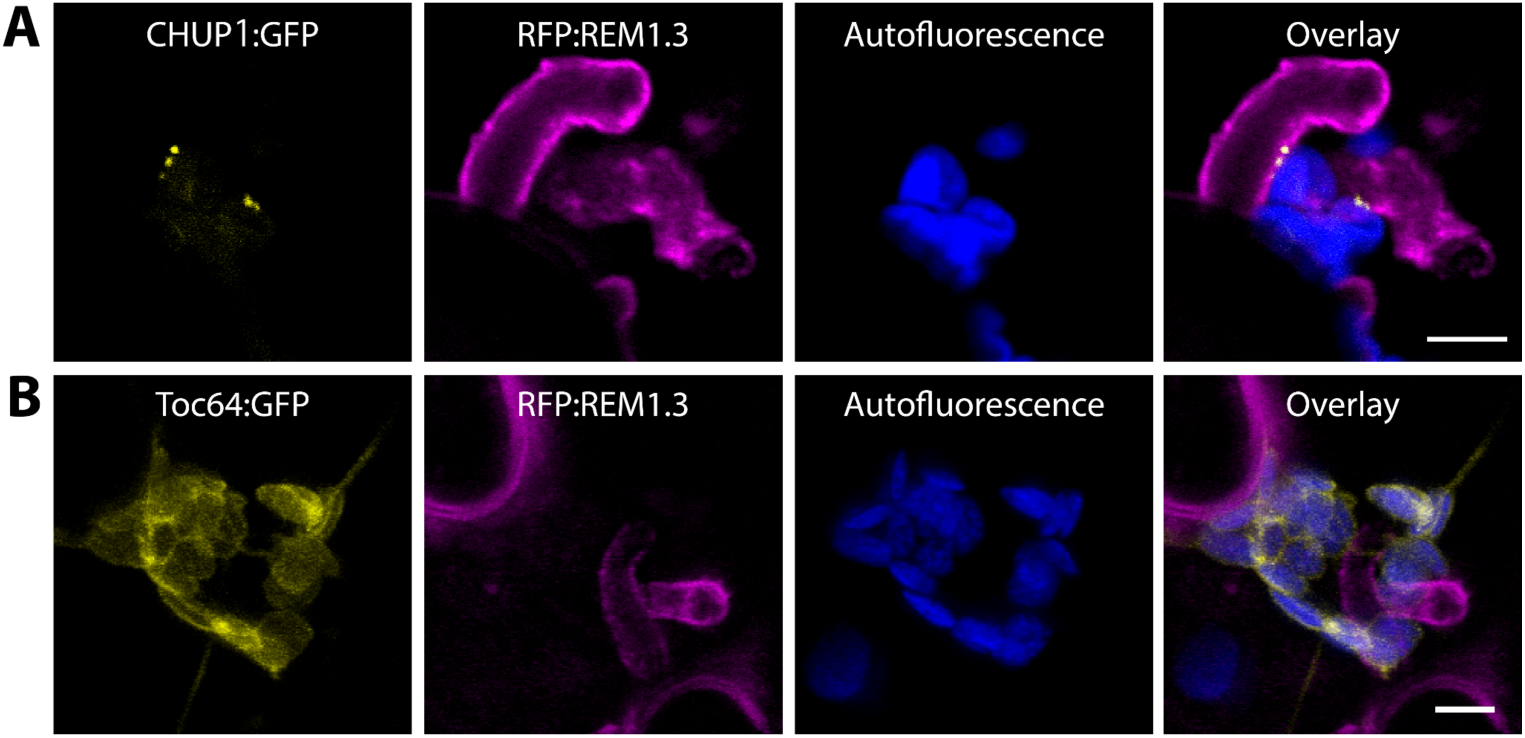
CHUP1 forms punctate accumulation at chloroplast-EHM MCS, but the control Toc64 does not. Additional representative images for Figures 2C and 2D. Confocal micrographs of *N. benthamiana* leaf epidermal cells transiently expressing either (A) CHUP1:GFP, or (B) Toc64:GFP, with RFP:REM1.3. The leaves were infected with WT *P. infestans* spores at 6 hpi. Images were taken at 3 dpi. REM1.3 is used as an EHM marker. Presented images are single plane images. Scale bars represent 5 µm.

**Figure S3.**
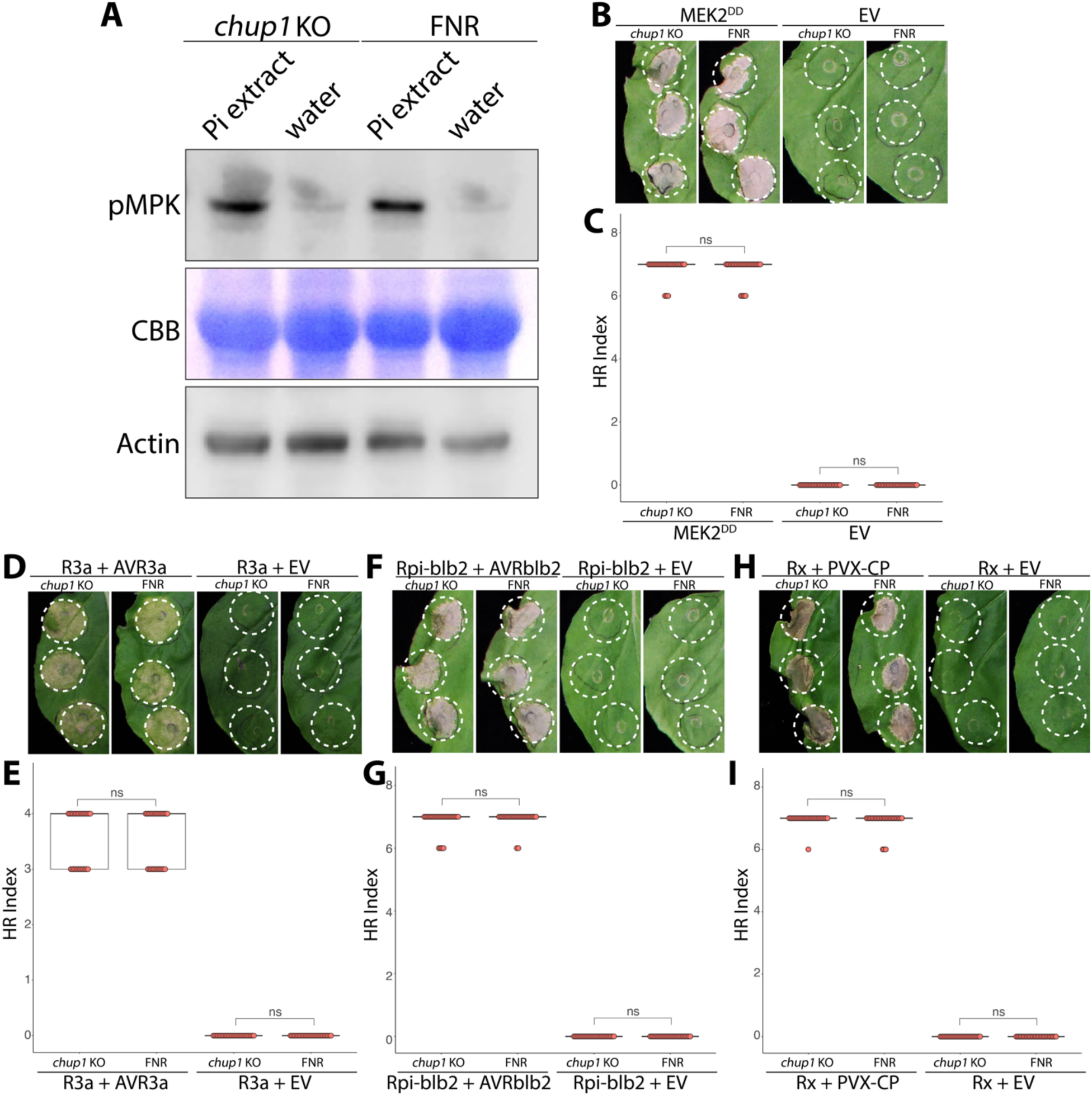
The activation of PAMP- or effector-triggered immunity is not impaired in the absence of CHUP1. (A) Western blot analysis of phosphorylated MAPKs in *chup1* KO and FNR plants after a 24-hour treatment with either Pi extract or water. Coomassie brilliant blue staining (CBB) of the gel post protein transfer and actin detection were performed for loading controls. (B-I) For all cell death assays, daylight images were taken, and cell death was scored at 3 dpi. Statistical significance was assessed using Mann-Whitney U test in R. ns = not significant. (B) Representative *chup1* KO and FNR *N. benthamiana* leaves infiltrated with MEK2^DD^ or EV control. (C) Dot plot illustrating that the differences in cell death induced by MEK2^DD^ in *chup1* KO (7, *n* = 27 spots) and FNR plants (7, *n* = 27 spots) are not significant. (D) Representative *chup1* KO and FNR *N. benthamiana* leaves infiltrated with R3a and AVR3a or EV control. (E) Dot plot illustrating that the differences in cell death induced by R3a with AVR3a in *chup1* KO (4, *n* = 27 spots) and FNR plants (4, *n* = 27 spots) are not significant. (F) Representative *chup1* KO and FNR *N. benthamiana* leaves infiltrated with Rpi-blb2 and AVRblb2 or EV control. (G) Dot plot illustrating that the differences in cell death induced by Rpi-blb2 with AVRblb2 in *chup1* KO (7, *n* = 27 spots) and FNR plants (7, *n* = 27 spots) are not significant. (H) Representative *chup1* KO and FNR *N. benthamiana* leaves infiltrated with Rx and PVX-CP or EV control. (I) Dot plot illustrating that the differences in cell death induced by Rx with PVX-CP in *chup1* KO (7, *n* = 27 spots) and FNR plants (7, *n* = 27 spots) are not significant.

**Figure S4.**
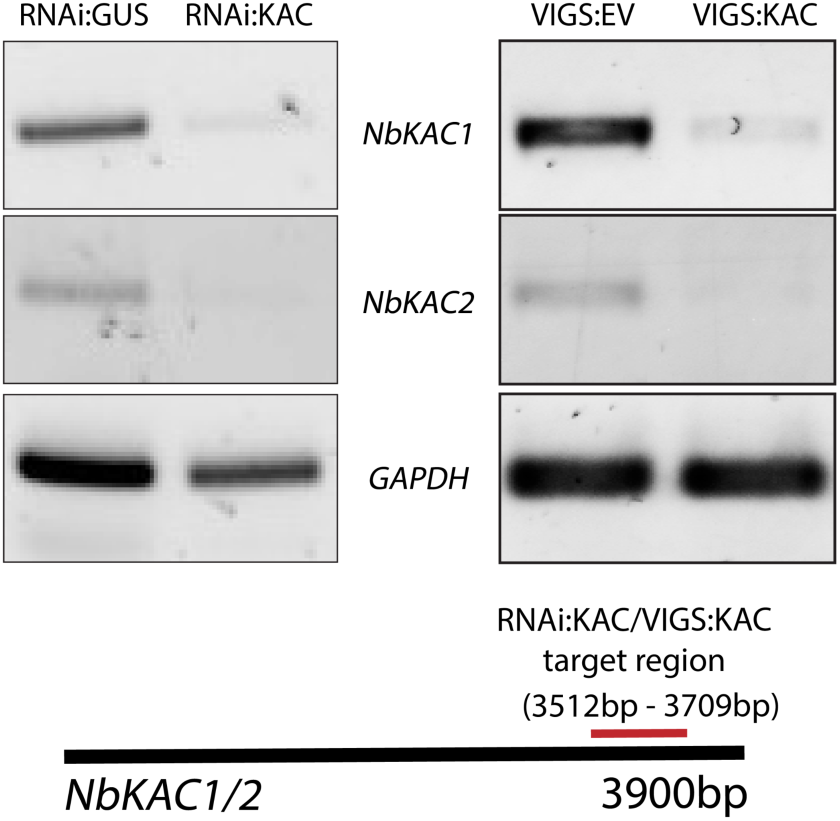
Validation of KAC1 silencing using the RNAi:KAC construct and VIGS:KAC plants. Hairpin plasmids (pRNAi-GG) targeting KAC1 or the GUS reporter gene were infiltrated into *N. benthamiana* leaves. The expression levels of the targeted genes were evaluated via RT-PCR at 3 days post-silencing (left panel). Constructs carrying TRV1 with TRV2-GG targeting *NbKAC1* and *NbKAC2* or the control TRV2:EV were infiltrated into *N. benthamiana leaves*, and the expression levels of the targeted genes were assessed via RT-PCR at 3 weeks post-VIGS (right panel). RT-PCR analysis utilized primers KAC1_RTPCR_F and KAC1_RTPCR_R for *NbKAC1*, and KAC2_RTPCR_F and KAC2_RTPCR_R for *NbKAC2*. The results confirmed gene silencing of *NbKAC1* and *NbKAC2* using the RNAi:KAC construct (left panel) and of the VIGS:KAC plants (right panel). Glyceraldehyde 3-phosphate dehydrogenase (GAPDH) served as the internal control, using primers GAPDH_RTPCR_F and GAPDH_RTPCR_R for assessment. The cDNA was synthesized from total RNA.

**Figure S5.**
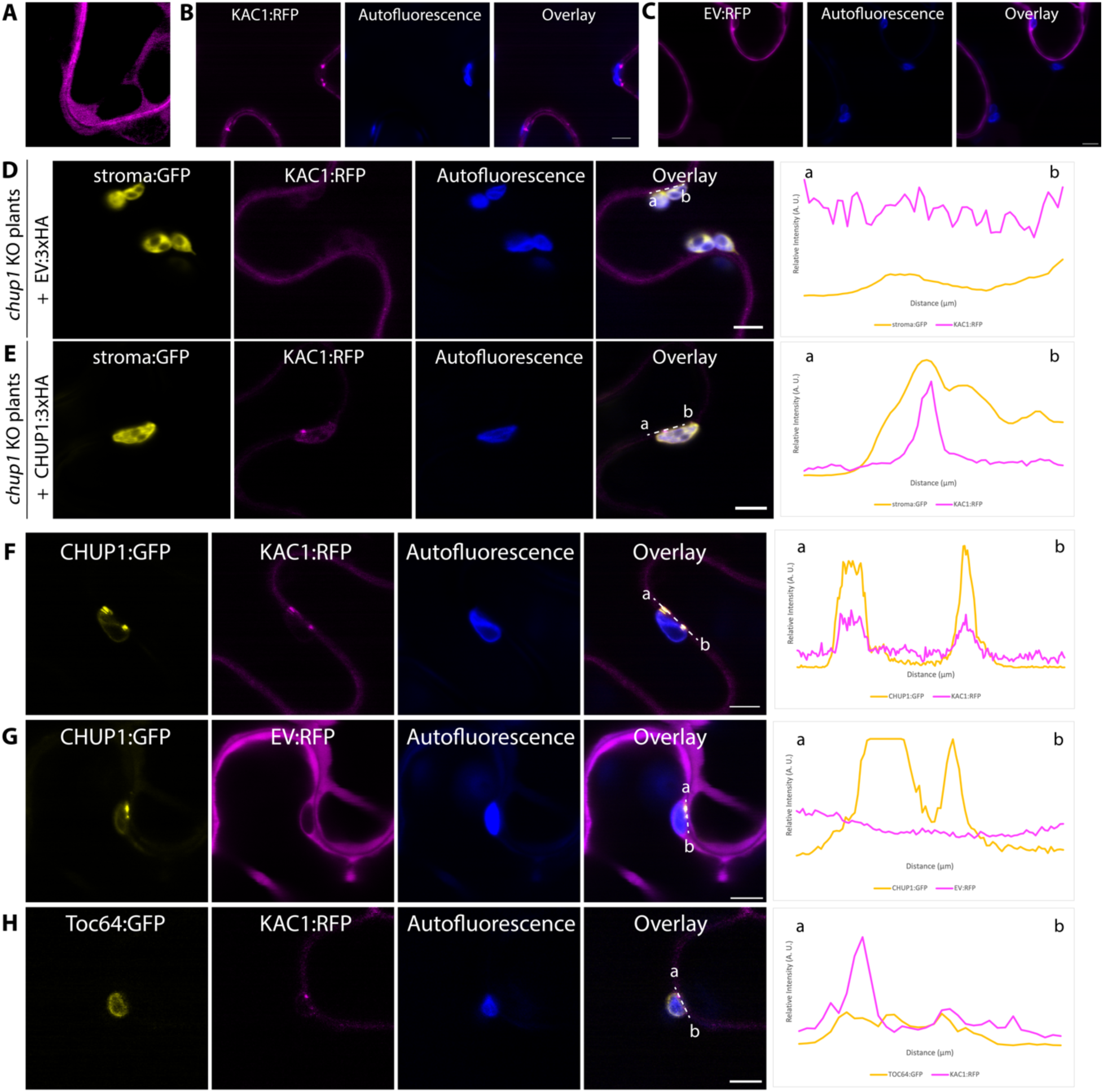
KAC1 interacts with CHUP1 and they colocalize at chloroplast-PM MCS. (A) KAC1:RFP localizes to the PM and cytosol. Confocal micrograph of *N. benthamiana* leaf epidermal cells transiently expressing KAC1:RFP. (B-C) KAC1:RFP forms punctate accumulation at chloroplast-PM MCS, but the control EV:RFP does not. Confocal micrographs of *N. benthamiana* leaf epidermal cells transiently expressing either (B) KAC1:RFP or (C) EV:RFP. (D-E) CHUP1 is required for KAC1 to form punctate structures at chloroplast-PM MCS. Confocal micrographs of *chup1* KO *N. benthamiana* leaf epidermal cells transiently expressing KAC1:RFP with (D) EV:3xHA, or (E) CHUP1:3xHA. GFP channel depicts chloroplast stroma in *chup1* KO plants. (F-H) CHUP1, but not Toc64, colocalizes with KAC1 at punctate structures at chloroplast-PM MCS. Confocal micrographs of *N. benthamiana* leaf epidermal cells transiently expressing (F) CHUP1:GFP and KAC1:RFP, (G) CHUP1:GFP and EV:RFP, and (H) Toc64:GFP and KAC1:RFP. All presented confocal images are single plane images. Images were taken at 3 dpi. Autofluorescence channel depicts chloroplasts. Transects in overlay panels correspond to line intensity plots depicting the relative fluorescence across the marked distance. Scale bars represent 5 µm.

**Figure S6.**
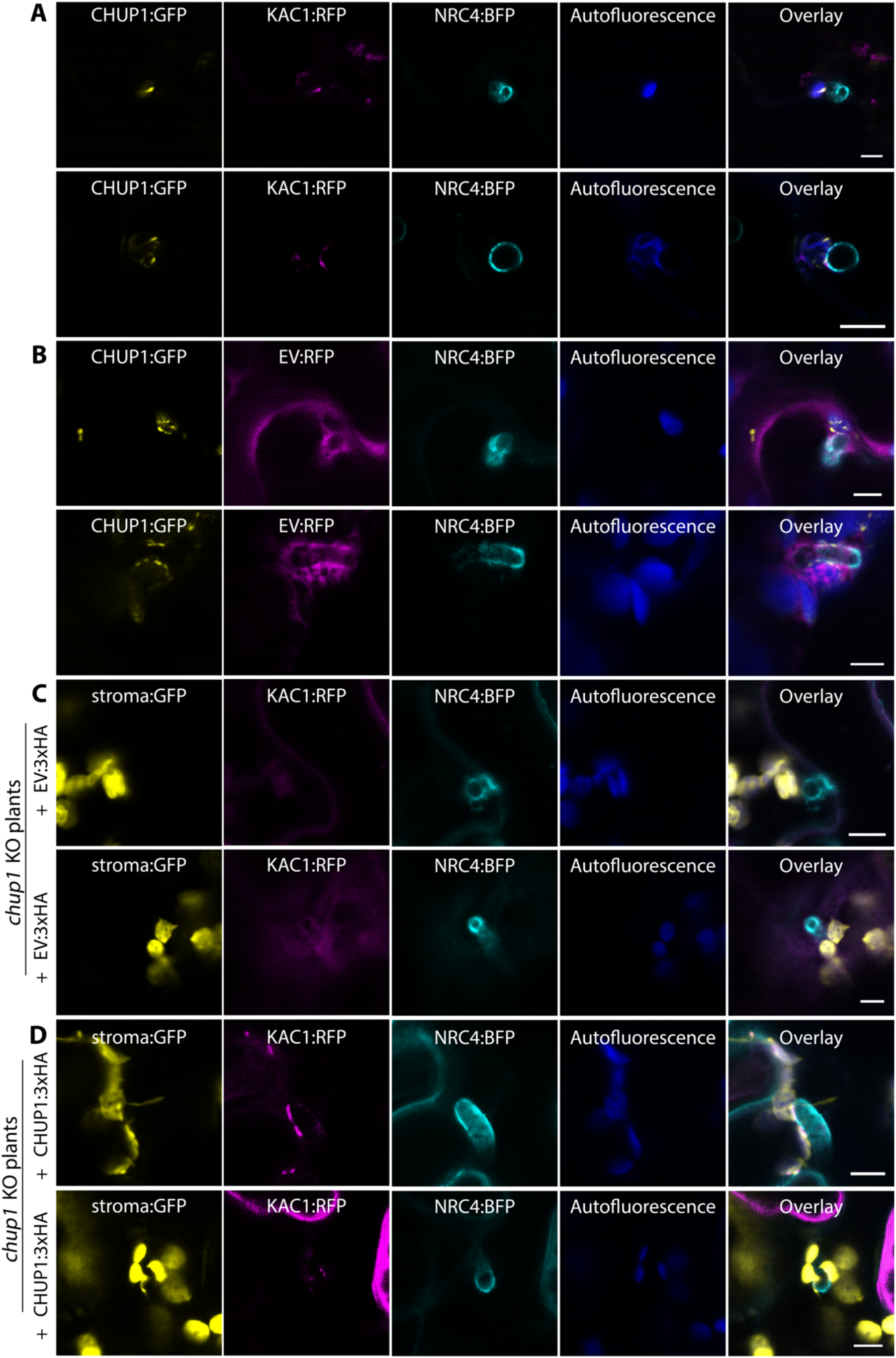
KAC1 colocalizes with CHUP1 at chloroplast-EHM MCS. Additional representative images for Figures 5A – 5D. (A-B) Additional representative images for Figures 5A and 5B. CHUP1 colocalizes with KAC1 at punctate structures at chloroplast-EHM MCS. Confocal micrographs of *N. benthamiana* leaf epidermal cells transiently expressing CHUP1:GFP and NRC4:BFP, with (A) KAC1:RFP, or (B) EV:RFP. NRC4:BFP acts as an EHM marker. The leaves were infected with WT *P. infestans* spores at 6 hpi, and imaged at 3 dpi. (C-D) Additional representative images for Figures 5C and 5D. CHUP1 is required for KAC1 to form punctate structures at chloroplast-EHM MCS. Confocal micrographs of *chup1* KO *N. benthamiana* leaf epidermal cells transiently expressing KAC1:RFP and NRC4:BFP with (C) EV:3xHA, or (D) CHUP1:3xHA. NRC4:BFP acts as an EHM marker. The leaves were infected with WT *P. infestans* spores at 6 hpi, and imaged at 3 dpi. All presented confocal images are single plane images. Autofluorescence channel depicts chloroplasts. GFP channel depicts chloroplast stroma in *chup1* KO plants. Scale bars represent 5 µm.

**Figure S7.**
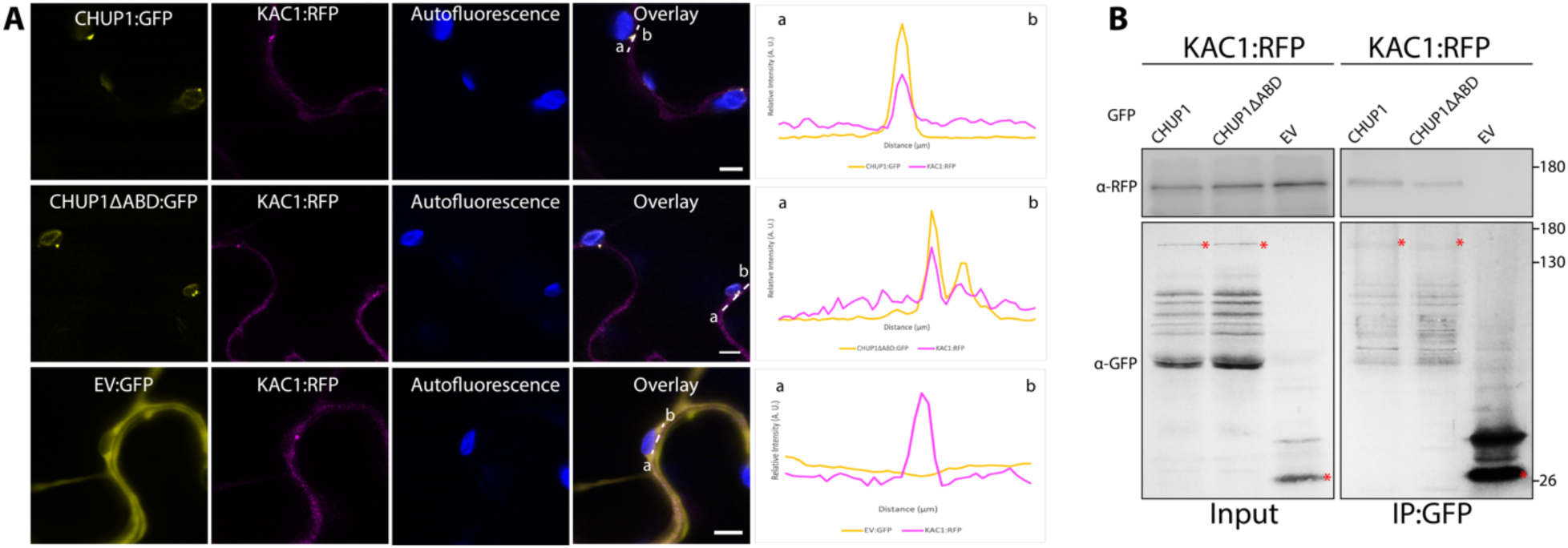
CHUP1 colocalizes and interacts with KAC1 independent of the actin-binding domain of CHUP1. (A) CHUP1 colocalizes with KAC1 at punctate structures at chloroplast-PM MCS independent of its actin-binding domain. Confocal micrographs of *N. benthamiana* leaf epidermal cells transiently expressing CHUP1:GFP and KAC1:RFP (1^st^ row), CHUP1ΔABD:GFP and KAC1:RFP (2^nd^ row), and EV:GFP and KAC1:RFP (3^rd^ row). All presented confocal images are single plane images. Images were taken at 3 dpi. Autofluorescence channel depicts chloroplasts. Transects in overlay panels correspond to line intensity plots depicting the relative fluorescence across the marked distance. Scale bars represent 5 µm. (B) CHUP1 interacts with KAC1 independent of its actin-binding domain. KAC1:RFP was transiently co-expressed with either CHUP1:GFP, CHUP1ΔABD:GFP, or EV:GFP. IPs were obtained with anti-GFP antibody. Total protein extracts were immunoblotted. Red asterisks indicate band sizes. Numbers on the right indicate kDa values.

**Table S1.**
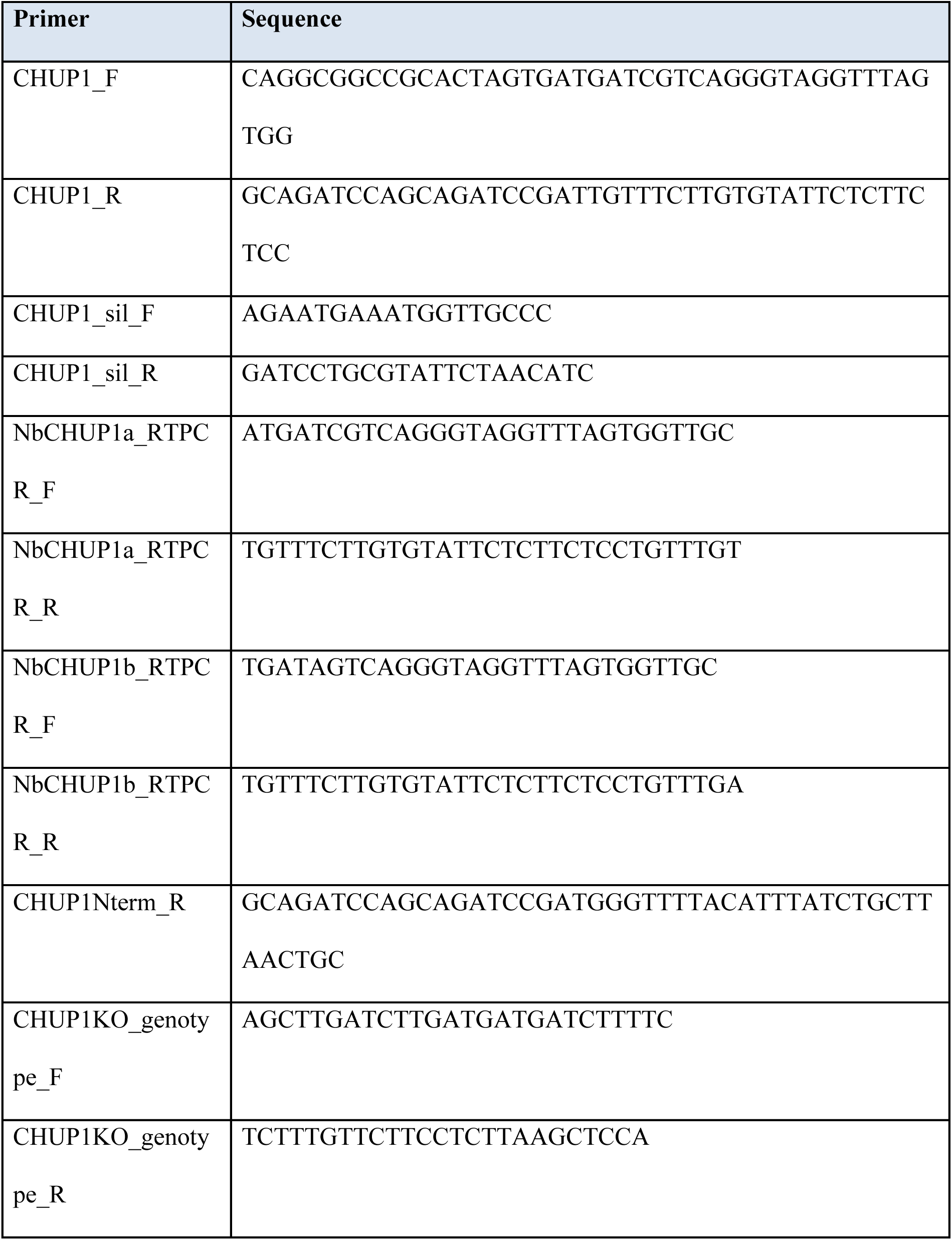
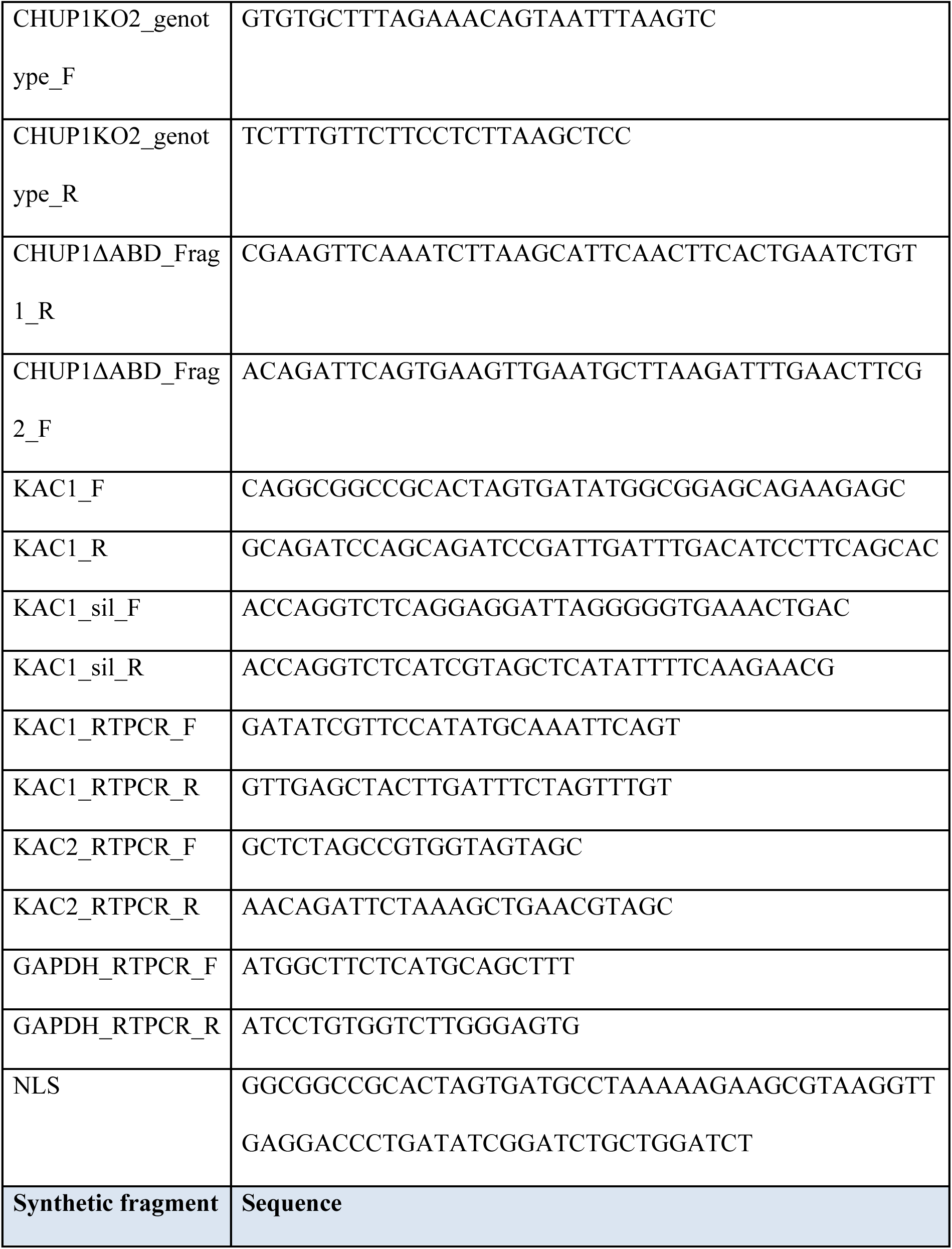
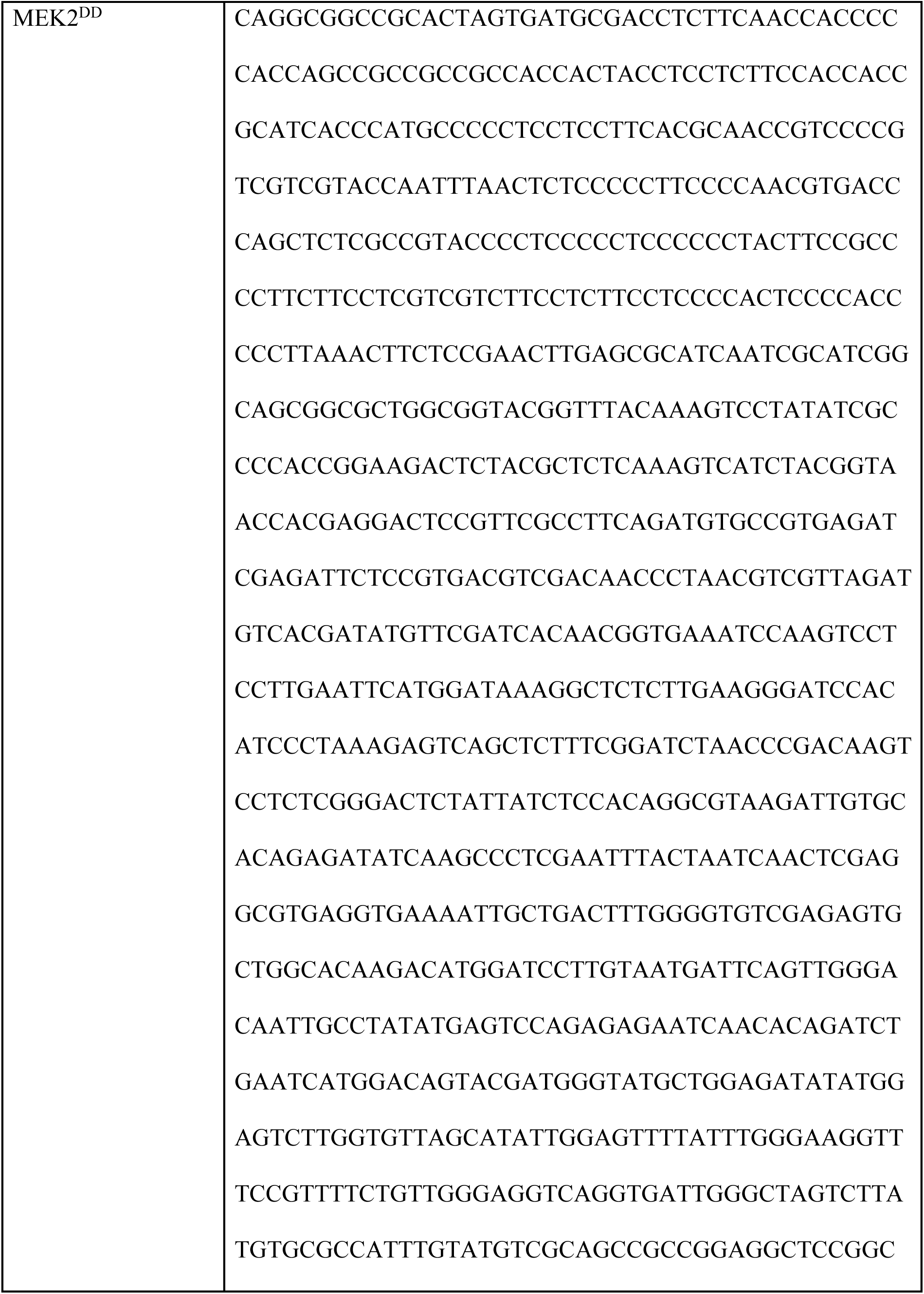
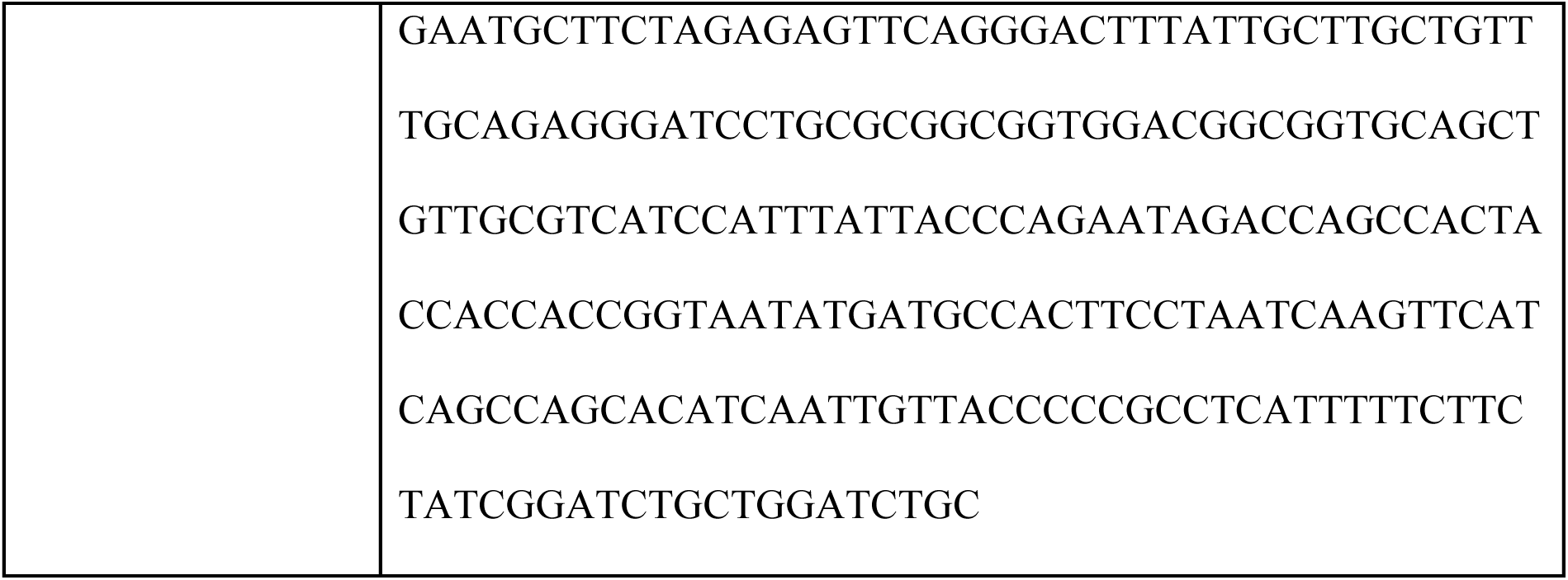
List of primers and synthetic fragment used in this study.

**Table S2.** List of constructs used in this study.

| Construct | Reference |
| --- | --- |
| Vector (pK7WGF2 domesticated for Gibson assembly) | 64 |
| Vector (pRNAI-GG) | 61 |
| Vector (TRV2-GG) | 60 |
| TRV1 | 65 |
| TRV2:EV | 60 |
| TRV2:KAC | This work (primers: KAC1_sil_F and KAC1_sil_R) |
| TRV2:CHUP1 | This work (primers: CHUP1_sil_F and CHUP1_sil_R) |
| RNAi:KAC | This work (primers: KAC1_sil_F and KAC1_sil_R) |
| RNAi:GUS | 66 |
| KAC1:RFP | This work (primers: KAC1_F and KAC1_R) |
| CHUP1:GFP | This work (primers: CHUP1_F and CHUP1_R) |
| CHUP1:3xHA | This work (primers: CHUP1_F and CHUP1_R) |
| CHUP1<sup>Nterm</sup>:GFP | This work (primers: CHUP1_F and CHUP1Nterm_R) |
| CHUP1ΔABD:GFP | This work (primers: CHUP1_F and CHUP1ΔABD_Frag1_R, CHUP1ΔABD_Frag2_F and CHUP1_R) |
| Toc64:GFP | 67 |
| RFP:REM1.3 | 42 |
| NRC4:BFP | 68 |
| EV:GFP | 68 |
| EV:RFP | 59 |
| 3xHA:EV | 69 |
| NLS:BFP | This work (primer: NLS) |
| MEK2<sup>DD</sup>:3xHA | This work (synthetic fragment: MEK2<sup>DD</sup>) |
| AVR3a | Provided by TSLSynBio |
| R3a | 70 |
| GFP:AVRblb2 | 1 |
| RFP:Rpi-blb2 | 1 |
| PVX-CP | 71 |
| Rx | 71 |

